# The anatomy of a phenological mismatch: interacting consumer demand and resource characteristics determine the consequences of mismatching

**DOI:** 10.1101/2020.12.22.423968

**Authors:** Luke R. Wilde, Josiah E. Simmons, Rose J. Swift, Nathan R. Senner

## Abstract

Climate change has caused shifts in seasonally recurring biological events and the temporal decoupling of consumer-resource pairs – i.e., phenological mismatching (hereafter, ‘mismatching’). Despite the hypothetical risk mismatching poses to consumers, it does not invariably lead to individual- or population-level effects. This may stem from how mismatches are typically defined, where an individual or population are ‘matched or mismatched’ based on the degree of asynchrony with a resource pulse. However, because both resource availability and consumer demands change over time, this categorical definition can obscure within- or among-individual fitness effects. We investigated the effects of resource characteristics on the growth, daily survival, and fledging rates of Hudsonian godwit (*Limosa haemastica*) chicks hatched near Beluga River, Alaska. To do this, we developed models to identify the effects of resource characteristics on individual- and population-level processes and determine how the strength of these effects change throughout a consumer’s early development. We found that at the individual-level, chick growth and survival improved following periods of higher invertebrate abundance but were increasingly dependent on the availability of larger prey as chicks aged. At the population level, seasonal fledging rates were best explained by a model including age-structured consumer demand. Our study suggests that modelling the effects of mismatching as a disrupted interaction between consumers and their resources provides a biological mechanism for how mismatching occurs and clarifies when it matters to individuals and populations. Given the variable responses to mismatching exhibited by consumer populations, such tools for predicting how populations may respond under future climatic conditions will be critical for conservation planning.

## Introduction

Shifts in the timing of recurring biological events (i.e., phenology) are among the best documented effects of climate change on living systems (Parmesan and Yohe 2003). Higher spring temperatures have led to earlier peaks in seasonal resources (e.g., invertebrate biomass; Pearce-Higgins et al., 2005; Tulp & Schekkerman, 2008). Slower rates of phenological advancement at upper trophic levels, however, mean that future climate conditions will likely lead to a greater decoupling of consumer-resource pairs (i.e., ‘mismatching’; Both & Visser, 2001; Both et al., 2009). Despite the theoretical risks imposed by climate-induced mismatching, mismatches do not invariably lead to reduced individual fitness (Dunn et al. 2011, Corkery et al. 2019) or negative demographic effects for populations (Visser et al. 2012, Reed et al. 2013, Keogan et al. 2020). The inconsistent effects of mismatching may be due to among-individual variation (Reed et al. 2013), although the underlying assumptions of existing mismatch models have also received scrutiny (Visser and Gienapp 2019). Recent studies have proposed improved methodologies for studying mismatches (Kharouba and Wolkovich 2020), but overcoming the empirical-theoretical disconnect in phenological studies may first require an improved mechanistic framework to help elucidate the degree to which mismatching occurs (Takimoto and Sato 2020).

The match-mismatch hypothesis presents mismatching as the disrupted interaction between consumer demands and resource availability (Cushing 1990). Most empirical studies categorize individuals or populations as either ‘matched or mismatched’ depending on the synchrony between the timing of a single life-history event and resource availability (Cushing 1974, Visser et al. 1998). Contrary to this categorization, both resource availability and consumer demands vary over time, and being ‘matched’ does not guarantee that consumers have sufficient food (Saalfeld et al. 2019, Keogan et al. 2020). Rather, changes to continuous resource characteristics like quantity (i.e., biomass) and quality (i.e., body size) directly affect consumer fitness, but the effects of these factors are rarely measured in phenological studies. Moreover, energetic demand changes throughout an individual’s life (Yang and Rudolf 2010), meaning that an individual’s sensitivity to resource availability is not constant but is instead likely age-structured (Dunn et al. 2011). Viewing mismatching simply as asynchrony in time, instead of as the disrupted interaction between consumer demand and resource availability, can obscure the cumulative effects of mismatching and mask population-level consequences (Yang and Rudolf 2010, Kerby et al. 2012). Although many conceptual models have been proposed to address this issue, a more robust methodology to model mismatching as the interaction of dynamic consumer demands and resources is still lacking (Chmura et al. 2019, Visser and Gienapp 2019).

Incorporating both age-structured consumer demand and multiple facets of resource availability into mismatch models likely requires a re-examination of our statistical concept of mismatching (Visser and Both 2005, Kellermann and van Riper 2015). Phenologies are generally modelled as frequency curves on a temporal axis (Fig. 1; Cushing, 1974; Visser et al., 1998), whereby a population’s degree of match with their resource is estimated as the difference in peak dates (i.e., date models) or proportion of overlapping area (i.e., overlap models). However, both date and overlap models have been criticized in the literature (Lindén, 2018; Ramakers et al., 2020). While date and overlap models agree if consumer and resource curves are symmetrical (Fig. 1a,b), date models can be biased when phenologies are skewed or multimodal, or in cases of low resource availability (Fig. 1c,d,e). Because overlap models account for the full interaction of consumer demand and resource availability posed in the match-mismatch hypothesis (Kerby et al. 2012), they may be better able to capture the mechanism of mismatching.

**Figure 1.**
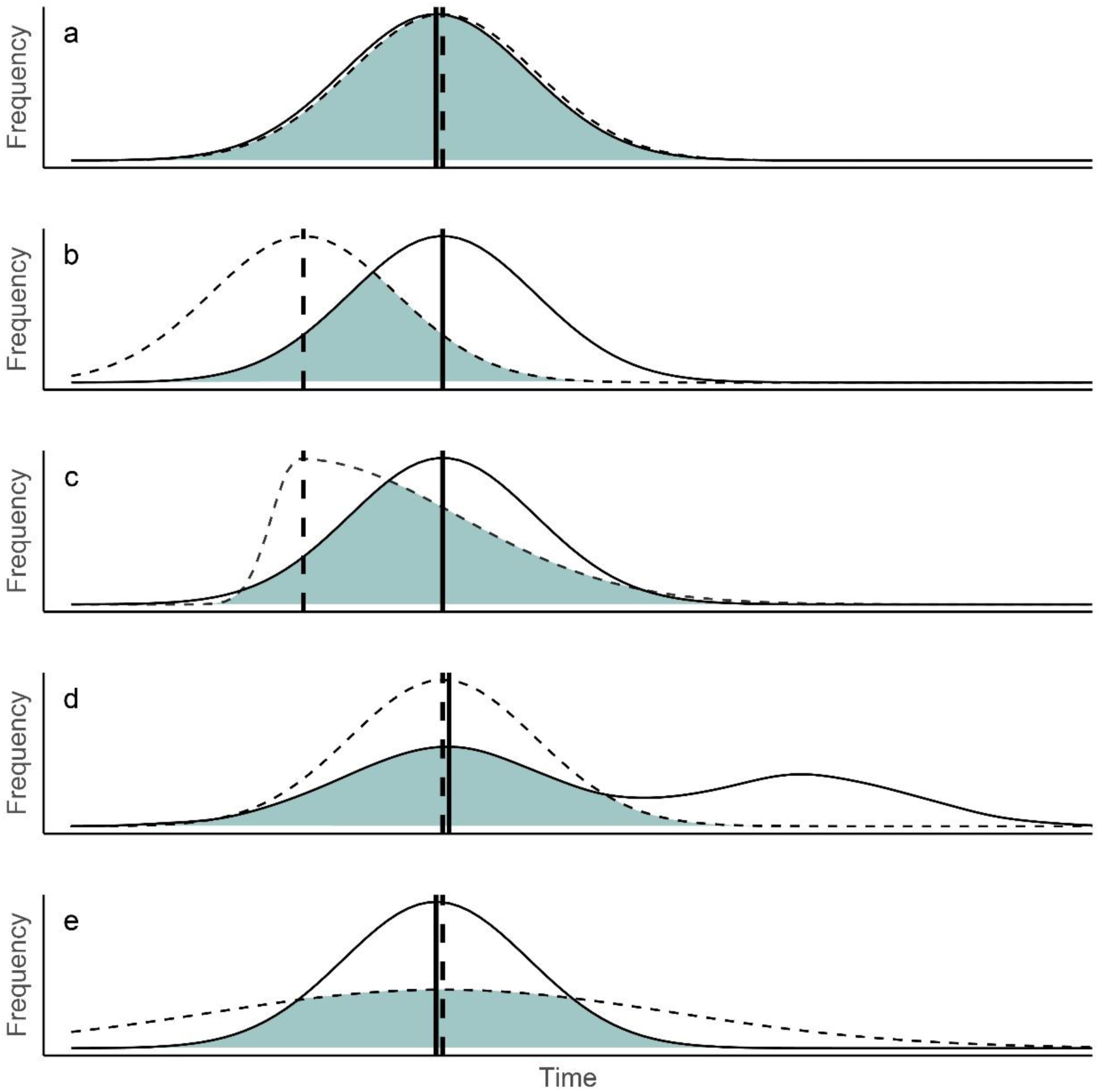
Peak dates (vertical lines) and frequency curves (phenologies) of consumers (solid) and resources (dashed). Difference in peak dates and peak overlap (shaded area; percent area under the curve) models are approximately equivalent when both the consumer (solid) and resource (dashed) curves are symmetrical (a, b). In this case, mismatching is a function of temporal displacement. However, date and overlap model estimates differ when either curve is skewed (c), the consumer phenology is multimodal (d), or the curves are aligned but have low overlapping area due to reduced resource abundance (e).

The inconsistent performance of overlap models may result from an inaccurate representation of consumer demand (Yang and Rudolf 2010, Kerby et al. 2012, Lindén 2018). Existing ‘peak demand’ overlap models estimate consumer demand from a single life-history event or timepoint in development, such as when individual growth rates are maximized (Fig. 2a; Leung et al., 2018). This approach, however, ignores demand prior to or following this peak, and results in a less realistic demand curve (Fig. 2b; Kerby et al., 2012; Lindén, 2018). Because animals require increasing energy as they develop, their sensitivity to the low resource availability associated with mismatching is likely to change over time. As a result, measuring the consequences of a mismatch from one timepoint could shroud cumulative effects (Yang and Rudolf 2010) and mask variation among individuals of differing ages (Reed et al. 2013). The growing availability of metabolic data and advances in survival analyses modeled in a Bayesian framework now allow for the direct simulation of the age- or stage-specific effects of mismatching. By modelling cumulative consumer demand as a function of the population age-structure, a ‘whole demand’ model incorporates the increasing metabolic demands of individuals as they age (Fig. 2c). As a result, the whole demand curve quantifies overlap at the demand curve’s upper tail when per-capita consumer demands are likely greatest (Fig. 2d; Kerby et al., 2012). Accurately modelling consumer demand and competing factors of resource availability may therefore be key to defining how mismatching occurs and when it should matter to populations.

**Figure 2.**
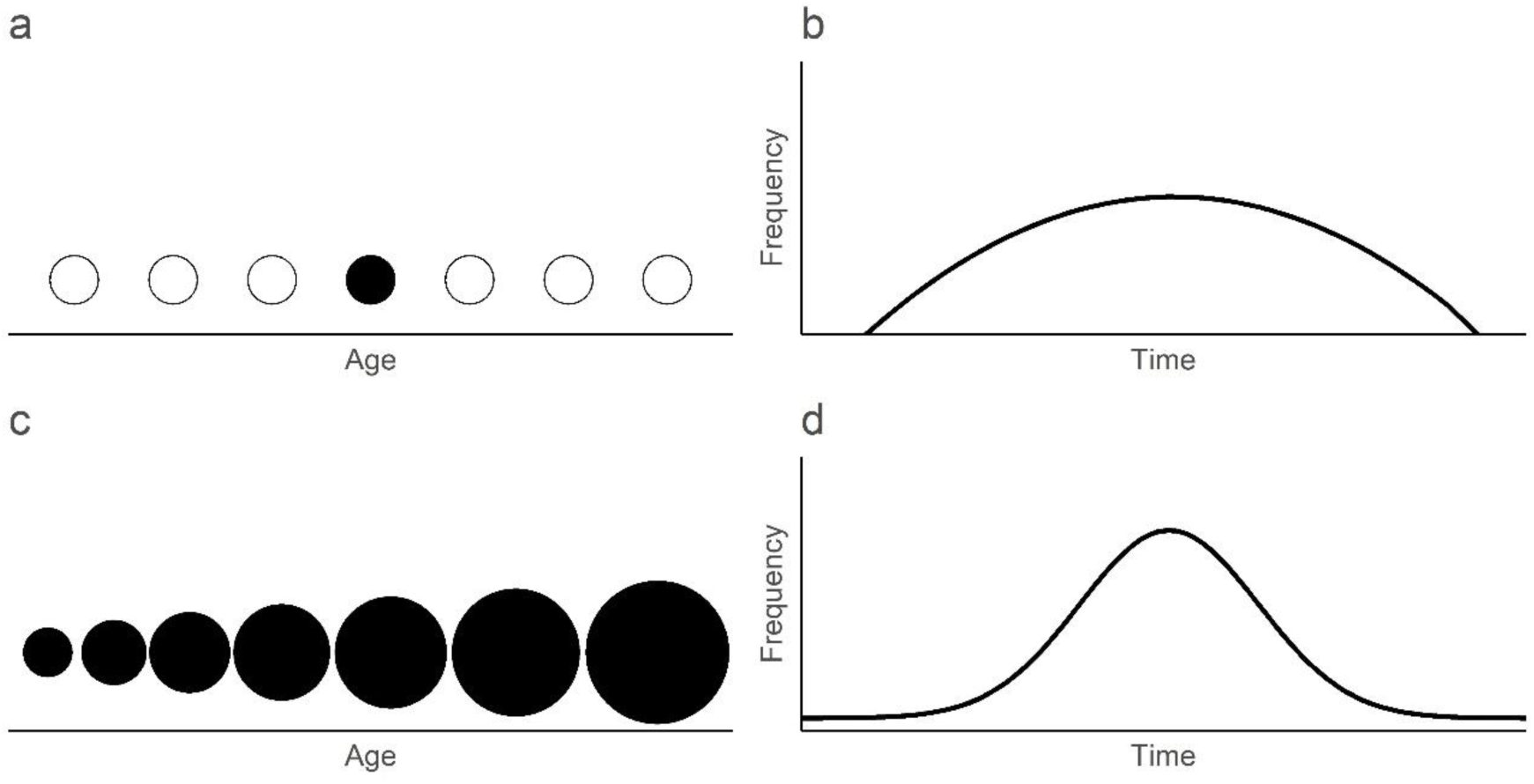
The peak demand model estimates consumer phenologies from the daily frequency of individuals at a single point in development (e.g., peak growth rate; a). Fitting a curve to pseudo-discrete data of this kind results in a simplified curve (b). However, since resource demand increases throughout development (c), including the cumulative demand of all individuals for each day of the season produces a curve with well-defined tails (d). Filled circles are time points in an individual’s development considered by the model. Circle size corresponds with hypothetical energy requirements at each timepoint. Curves are from predictions from a generalized additive mixed model (GAMM) performed on data collected in our study (see Methods and Results).

Migratory birds provide a powerful avenue to re-examine the effects of mismatches under this new framework. Long-distance migrants represent some of the canonical examples of mismatches because of their use of endogenous cues to time migrations and reproduction (Both and Visser 2001), and their reliance on seasonal resource pulses to achieve rapid offspring growth (Schekkerman and Visser 2001). While many studies have identified individual-level fitness effects resulting from mismatches, few have found corresponding population-level consequences (Visser and Both 2005, Dunn et al. 2011). Hudsonian Godwits (*Limosa haemastica*; hereafter, ‘godwits’) are a case-in-point. Godwits breed in three disjunct populations spread across the Nearctic (Walker et al. 2011). Like other shorebird species (Kwon et al., 2019), godwits breeding in Alaska have kept pace with recent phenological changes in peak resource availability while those breeding in Hudson Bay have not (Senner 2012). Despite mismatches affecting the survival of individual godwit chicks in Hudson Bay, there have been few apparent population-level consequences (Senner et al. 2017). Furthermore, much of the interannual variation in the fledging rates of Alaskan godwits is not explained by predation or density-dependent processes (Senner et al., 2017; Swift et al., 2017a, 2018; Wilde et al., in revision). The observed interannual variation may instead result from a potential correlation between early snowmelt and low seasonal godwit fledging rates, suggesting that mismatching may be occurring and having demographic consequences (Saalfeld et al. 2019).

Updating our conceptualization of mismatches may document the effects of resource availability on consumer fitness that our previous attempts based on the categorical view of mismatching have missed. Therefore, we investigated how dynamic consumer demand and resource characteristics interact to influence the potential for mismatching in the Alaskan population of godwits. We developed mechanistic models that integrate metabolic and resource availability information at the individual- and population-levels. We first explored how the timing, abundance, and quality of resources have changed over the course of the breeding season. Then, we investigated the effects of invertebrate abundance and size on the growth and survival of godwit chicks. We hypothesized that mismatching affects individual fitness differently throughout development and predicted that chick growth and survival would improve with more abundant and larger prey, with the ‘size’ effect increasing with age. Lastly, we investigated the influence mismatching has on godwit population dynamics. We hypothesized that mismatching in godwits is simultaneously a function of both consumer demand and resources. We therefore predicted that the whole demand model would explain population-level effects better than alternatives. Identifying how resources interact with consumer demands will provide evidence for the mechanism underlying mismatches and help better connect mismatching to demographic processes.

## Methods

### Study area and godwit chick monitoring

During 2009–2011, 2014–2016, and 2019, we monitored godwits on two plots – North (550 Ha) and South (120 Ha) – near Beluga River, Alaska (61.21°N, 151.03°W; hereafter, ‘Beluga’; Appendix S1: Fig. S1). Both plots consist of freshwater ponds and black spruce outcroppings (*Picea mariana*) dominated by dwarf shrub and graminoids surrounded by boreal forest (Swift et al. 2017a, 2017b).

Each season (early-May to mid-July: 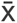 = 78 days), we censused both plots for godwit nests, locating, on average, 23 nests per season (range: 11–33). For each nest, we estimated hatch date by floating the eggs to estimate the age of the nest (Liebezeit et al. 2007). We monitored the nests’ survival every 2–3 days until eggs showed starring or pipping, after which, we monitored them daily until hatch. We captured newly hatched chicks and collected morphometric measurements and body weights on all chicks in the brood. We uniquely marked each with a leg-flag and U.S. Geological Survey metal band, neither of which are likely to impact survival (Sharpe et al. 2009). Some nests hatched before detection each season (range: 0–4). We opportunistically captured chicks from these broods off-nest and estimated their age from weight measurements. Because we included chicks captured off-nest, we are confident that we found all broods each season given the small size of the study area and the conspicuousness of godwit broods.

We monitored the survival of 1–2 chicks chosen randomly from each brood (range = 7– 23 chicks per season), except for one brood in 2019 from which we captured two chicks > 14 days after their estimated hatch date. We attached a 0.62 g very high frequency radio transmitter (Holohil Systems Ltd.) above the uropygial gland by clipping the feathers and using skin-safe, cyanoacrylate glue to attach the radios directly to the skin. We relocated each radioed chick every 2–3 days by walking the entirety of the study plot and recording telemetry azimuths from <100-m of the tending parent(s)’s location. When no parents were present, we scanned for signals in all directions from a chick’s last known location. We attempted to recapture radioed chicks weekly to reapply glue and measure their body mass to the nearest gram, minus the transmitter weight, using a magnetic field scale.

Godwits are fully flight capable, or ‘fledged’, after ∼28 days (Walker et al. 2011). However, because our radios had an expected lifespan of 21-days (range: 17–30 d; Holohil Systems, 2021), we considered chicks that survived to 21 days to have fledged. We confirmed mortalities when possible and assumed that chicks had died after three consecutive failed location attempts. While it is possible that some radios failed, we did not record any instances in which this occurred. Nonetheless, we are confident that the lifespan of our radios was sufficient for capturing survival for two reasons: first, the chicks that died during monitoring did so on average, by 3.6 days of age (SD: 8.3; range: 0–16), and second, we never resighted a chick that was presumed dead during weekly censuses of the bog and nearby foraging areas (N.R. Senner, University of South Carolina, unpublished data, 2019).

### Resource monitoring

During 2009–2011, 2012, 2014–2016, 2017, and 2019, we monitored the abundance and body size of invertebrates. In 2009–2011, 2014–2016, and 2019, we monitored invertebrates for an average of 67 days (range = 61–78) simultaneous with our godwit monitoring. Additionally, we monitored invertebrates, but not godwit nests, for 38 and 5 days in 2012 and 2017, respectively, as these were shortened seasons with limited crews. Passive traps are a good proxy of resource availability to foraging shorebird chicks (McKinnon et al. 2012). We thus collected invertebrates each day along two, 50-m transects consisting of five traps placed within mesic godwit breeding habitat (Brown et al. 2014, Senner et al. 2017). We used two trap styles: pitfall traps (10 × 15 cm) filled with 10 cm of 75% ethanol from 2009–2012, and modified malaise traps (see Leung et al., 2018) filled with 3 cm of 75% ethanol from 2014–2019. We cleared and replenished traps every 24 hours.

We identified invertebrates to Order and measured body-lengths to the nearest 0.5-mm. We converted lengths to inferred dry mass using published, taxon specific length-weight relationships (Ganihar, 1997; Rogers et al., 1977).

### Statistical Analyses

#### Interannual resource variation

To examine resource availability over the course of our study, we investigated how the (1) date of the seasonal peak, (2) daily biomass (transect^-1^ day^-1^; dry mass, mg), and (3) daily median invertebrate body size (per-capita dry mass, mg) changed across seasons. Because godwit chicks are gape limited and rarely consume larval invertebrates, we restricted our analysis to only include individual invertebrates that are potential prey for godwits by subsetting our data to adult invertebrates with lengths of 1.5–9 mm (Schekkerman and Boele 2009). We also excluded the shortened 2012 and 2017 seasons from our analyses of seasonal peak timing but included them in tests of daily biomass and daily median body size. We treated the transects as replicates, averaging the total daily biomass collected along each transect for each day. We then estimated overall and order-specific seasonal peaks using the first derivative of quadratic curves (Julian day + Julian day^2^) fit to the daily biomass within each season. We built separate mixed-effect models to estimate linear trends over time in the overall and order-specific dates of peak abundance (i.e., peak dates), daily biomass, and daily median body size. We included a random intercept for ‘trap type’ in all our models using the *lmer* function (package ‘lme4’, Bates et al. 2015) in the R programming environment (v4.0.3, R Core Team 2020). In the daily biomass and daily median invertebrate body size models, we also included a random intercept of sample date to compare trends within days of the season over the course of the study.

To identify potential changes in the composition of the invertebrate assemblage, we repeated the above analyses with each of the six Orders that comprised 91.6% of all observed invertebrates – Araneae (20.5%), Hymenoptera (18.4%), Coleoptera (17.5%), Diptera (16.2%), Acari (11.3%), and Hemiptera (7.7%; Appendix S1: Fig. S2). We excluded Collembola (8.3%) from our analyses, as they are primarily aquatic and thus unlikely prey for godwit chicks and were also poorly recorded, due to their low frequency from 2009–2012. We standardized response variables by dividing by two standard deviation according to Gelman (2008), but report coefficients in their original units throughout the text. We considered response variables whose 95% confidence intervals did not include zero as biologically relevant.

#### Chick growth and body condition

We modelled chick growth with a logistic growth function using the ‘nlme’ package (Pinheiro et al. 2020) to predict the age-specific mass of chicks (Senner et al., 2017). Although godwit chicks may be sexually dimorphic (Loonstra et al. 2018), we lacked data on each individual’s sex and therefore pooled the sexes in our analyses. We set the asymptotic mass to the population’s mean adult mass (249 g; Senner et al., 2017). Next, we developed separate growth models with chick ID as a random intercept, and constant or seasonal growth coefficients and inflection points (Pinheiro and Bates 2000). We performed 100 iterations for each model and included site-specific estimates from Senner et al. (2017) as starting values. We compared 12 candidate models using Akaike’s Information Criterion scores corrected for small sample sizes (AIC_c_; Burnham & Anderson, 2002). We used candidate models with < 2 ΔAIC_c_ to estimate the predicted chick growth curve. Next, we calculated the body condition index (BCI) for each recaptured individual by dividing the observed weight gain since last capture by the curve-predicted weight gain over the same time.

To investigate whether and over what timescale resource characteristics influenced chick growth, we modelled BCI in relation to resource abundance and quality in all seasons with godwit monitoring except 2014, which lacked sufficient recaptures. We built a global, generalized additive mixed model (GAMM) with a gaussian error term that included (1) daily invertebrate biomass, (2) daily median invertebrate body size, and (3) hatch date as fixed effects (package ‘gamlss’; Rigby and Stasinopoulos 2005). We included random intercepts for (4) season and (5) brood ID. Lastly, we (6) smoothed the effect of chick age using a cubic spline to account for irregular sampling (every 7-days) throughout development. We first ensured that all additive covariates included in the same model showed minimal collinearity with a pairwise Pearson’s correlation coefficient (r) below 0.7 (Appendix S1: Table S1). Because resource abundance and quality could have either an immediate or cumulative effect, we determined the timescale over which these predictors influenced BCI: day of recapture or 1-day, 3-day, or 7-day averages, and used the timescale with the lowest AIC_c_ score in further analyses. We also determined whether including random intercepts – individual, brood, or season – improved model fit with an AIC_c_ comparison. Finally, we built a global model with the timescale and random effects that optimized fit and compared all candidate models using AIC_c_ scores. When no model had a model weight (*w_i_*) > 0.90, we used model averaging (ΔAIC_c_ < 4) within the ‘MuMIn’ package and report conditional average coefficients (Bartoń 2015).

#### Effect of resources on survival: constant or age-varying?

To determine how invertebrate biomass or body size affected daily chick survival, we built a Bayesian hierarchical survival model. We constructed daily encounter histories for all individuals, beginning with an individual’s hatch date and ending with their expected fledging date. Because we assumed the chicks that we could not relocate for three consecutive days were dead, we included two days of unknown fate to allow for Markov chain Monte Carlo (MCMC) prediction. We modelled encounter histories as a Bernoulli variable and assumed fates were known.

In the second portion of our model, we incorporated parameters hypothesized to influence chick survival. We constructed a logit-linear mixed model to estimate the additive effects of (1) daily invertebrate biomass, (2) daily median invertebrate body size, (3) hatch date, and (4) chick age, along with random intercepts for each (5) brood ID, (6) season, and (7) study plot. To reduce the influence of outlier values of daily biomass or daily median size measurements, we averaged our continuous parameters across 3-day periods (i.e., our relocation interval) and standardized all variables by dividing by two standard deviations (Gelman 2008). To test whether the effects of daily invertebrate body size or daily biomass varied with chick age, we built separate models with interactive terms between chick age and either daily median invertebrate body size or daily invertebrate biomass. Given their rapid growth rates, age is a proxy for metabolic rate in shorebird chicks (Schekkerman and Visser 2001, Williams et al. 2007). We compared these age-interaction models using deviance information criterion (DIC) and included the interaction from the model with the lower DIC score in all further models. We chose diffuse priors for all our predictors (Normal(0, τ)) and constrained random intercepts close to 0 (mean = N(0, 1000), SD = Uniform(0, 25)).

To identify the top model, we performed model selection using the indicator-variable approach (Link and Barker 2006, Converse et al. 2013). We again checked for collinearity between additive covariates with a pairwise Pearson’s correlation coefficient (Appendix S1: Table S2). Next, we assigned a Bernoulli variable (weights) with a 0.5 prior to each predictor to model its inclusion (1) or absence (0) from each MCMC sample. We maintained an equal number of parameters across samples by fixing the model variance, τ = *K* * Gamma(3.29, 7.8), for all parameters, where *K* is the number of parameters (Link and Barker 2006). The posterior mean of the weight indicator is evidence for inclusion in the model. We calculated Bayes factors (BF) from predictor weights (Link and Barker 2006) and included predictors with BF > 3 in our top model along with their random intercepts. If an interaction term was selected for the top model, we included both additive terms included in the interaction regardless of their independent selection.

We constructed models of daily chick survival using the ‘runjags’ and ‘rjags’ packages (Appendix S2: R code S1; JAGS 4.1.0; Plummer, 2012, 2013; Denwood, 2016). Our models accessed three parallel chains to perform 5,000 iterations. We removed 600 and 1,000 iterations for adaptation and burn-in, respectively, with a one-third thinning factor. We assessed model performance based on the values of the Gelman-Rubin statistic < 1.1 and chain mixing (Gelman and Rubin 1992). For all tests, we report the beta coefficients in logit-form, 95% credible interval, and Bayesian p-value (probability of slope ≠ 0).

#### Population match and reproductive success

To quantify population-level mismatching, we built resource and consumer demand curves for each season. Additionally, we built competing demand curves from the ‘peak demand’ and ‘whole demand’ conceptual models (Fig. 2) to quantify the effect of dynamic consumer demand on godwit reproductive success. (1) *Peak demand*: Following Kwon et al. (2019), we calculated the number of all hatched godwit chicks expected to be 11-days old (i.e., age of peak growth rate; Senner et al. 2017) for each day of the season. We then converted both the daily values of invertebrate biomass (hereafter, ‘resource curve’) and counts of 11-day old chicks to their seasonal proportions. (2) *Whole demand*: We multiplied the maximum number of chicks of each age per day of the season by age-specific estimates of resting metabolic rate in godwit chicks (Williams et al. 2007). Resting metabolic rate approximates the amount of energy individuals use to maintain homeostasis and therefore represents an individual’s minimum energetic requirement independent of other factors (i.e., thermal environment). We then estimated the cumulative energetic requirements (Kilojoules d^-1^) of all chicks per day of the season and converted these to seasonal proportions to produce the whole demand curve.

We modelled the shape of the peak demand, whole demand, and resource curves using separate GAMs with a quadratic time function – Julian day + Julian day^2^ (Appendix S2: R code S2; Kwon et al. 2019). We restricted the analyses to 10 May – 10 July for comparison among seasons, which otherwise differed in length (Appendix S1: Table S3). We approximated error terms as a gaussian distribution (∼N[µ,σ]) and zero-inflated beta distributions (∼ zBeta[z|α, β]) for the peak demand and whole demand curves, respectively, and a beta distribution (∼Beta[α, β]) for the resource curve, all with logit-link functions. We fit the resource curve with a penalized spline (k = 10) to estimate mean predicted values for each day of the season while capturing the modality of the resource curve (Vatka et al. 2016). We then estimated the degree of overlap between the peak demand or whole demand curves and the resource curve by calculating the proportional area overlap using the *integrate.xy* function (‘sfsmisc’, Maechler 2020). We also estimated the (3) ‘curve height’ in each season (i.e., cumulative resource availability) from the area under the resource curve. Lastly, we calculated the (4) ‘difference in peak dates’ (i.e., synchrony) between the resource and peak demand curves in each season from the point at which each curve’s derivative was zero.

To determine how mismatching affected godwit reproductive success, we built four univariate linear models relating the different measures of mismatching to fledging rates – (1) peak demand, (2) whole demand, (3) difference in peak dates, and (4) curve height. We extrapolated daily survival rate estimates from our global Bayesian model to 28 days with the associated error using the Delta method (Powell 2007). We compared among the four models by calculating (1) model weights from their AIC_c_ scores and (2) the proportion of the variation in fledging rates they explained.

## Results

We located 142 godwit nests from 2009–2019, of which 128 survived to hatch. We individually marked 349 chicks (2009–2011, *n* = 195; 2014–2016, *n* = 106; 2019, *n* = 48) and attached radios to 128 chicks from 102 distinct broods. On average, radioed chicks survived to 9.4 days (SD: 8.4 d, range: 0–21). We relocated radioed chicks an average of 4.3 times (SD = 2.71, range: 1–19; n = 778), and we recaptured them 1.5 times (SD: 0.83; *n* = 103). In most cases of chick death, we located a carcass (38%) or found a detached radio in habitats clearly suggestive of predators (20%; e.g., on gull nesting island) within 2 days (range: 0–4 days) of the first failed relocation attempt.

### Interannual changes in resources

We recorded the body-lengths of 69,598 adult invertebrates across 14 orders, 41,298 of which were potential godwit prey (i.e., 1.5–9 mm in length) and thus considered in our analyses. Sample days showed wide variation in the biomass (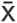 = 132.9 mg, range: 0–948.4 mg) and daily median body size (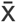 = 1.5 mg, range: 0.2–13.4 mg). We found no change in the timing of the predicted peak dates of all invertebrates (β = -1.68 ± 3.08 d, 95% CI = -3.34, 5.50 d; R^2^_m_ = -0.02; R^2^_c_ = -0.05) or among the individual orders over the course of the study (Fig. 3, left). However, both daily invertebrate biomass (β = -2.49 ± 0.50 mg, 95% CI = -3.49, -1.51; R^2^_m_ = -0.18; R^2^_c_ = -0.35; Fig. 3, center) and daily median invertebrate body size (β = -0.33 ± 0.03 mg, 95% CI = - 0.028, -0.37; R^2^_m_ = -0.13; R^2^ = -0.26; Fig. 3, right) did decrease over time at a rate of -2% and - 5% per year, respectively. At the order level, only Acari became more abundant over time (β = 0.20 ± 0.02 mg, 95% CI = 0.15, 0.25; R^2^_m_ = 0.14; R^2^_c_ = 0.38), while all other taxa became less abundant (Fig. 3). Additionally, Araneae (β = -0.67 ± 0.09 mg, 95% CI = -0.49,-0.85; R^2^_m_ = - 0.19; R^2^ = -0.28), Diptera (β = -0.24 ± 0.02 mg, 95% CI = -0.20, -0.29; R^2^ = -0.03; R^2^ = - 0.15), and Hemiptera (β = -0.23 ± 0.06 mg, 95% CI = -0.10, -0.35; R^2^_m_ = -0.11; R^2^ = -0.16) showed consistent decreases in body size over the course of the study (Fig. 3).

**Figure 3.**
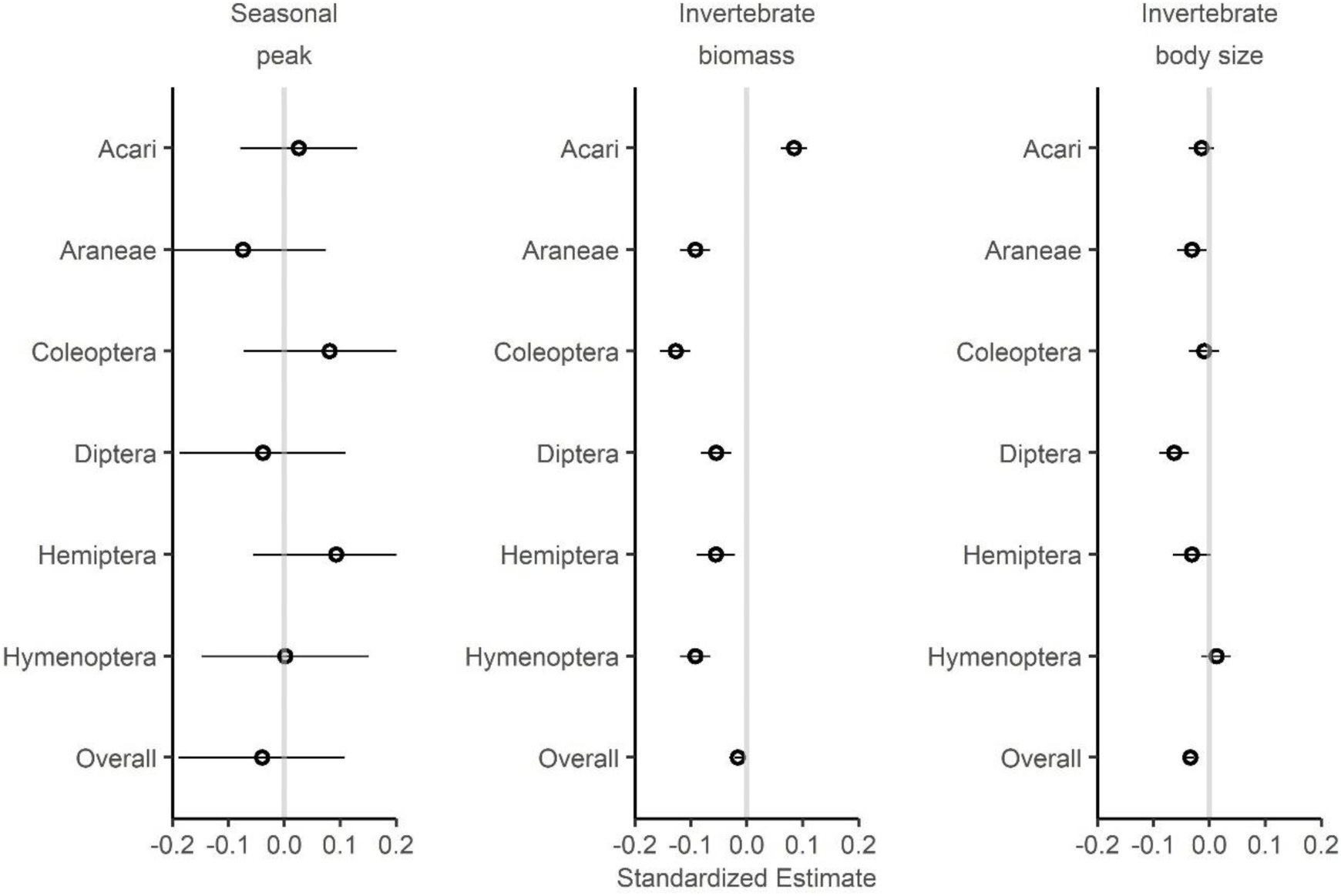
Interannual changes of within season peak timing (left), observed daily invertebrate biomass (center), and daily median invertebrate body size (right) of six common Orders and the invertebrate assemblage overall. Linear regression estimates are shown as hollow circles, with 95% confidence intervals shown as horizontal lines. Variables with no consistent effect had intervals that crossed zero (grey line).

We found opposing trends in daily biomass during the early and late portions of the godwit breeding season. Days during the nest incubation period (16 May–6 Jun) from 2014– 2019 had 83% higher invertebrate biomass than those from 2009–2012, but 41% lower biomass on days during the chick-rearing period (6 Jun – 4 Jul). Meanwhile, invertebrate body size was 42–72% smaller in the later period.

### Chick growth and body condition

We modeled godwit chick growth from 103 mass-at-capture measurements taken following the initial measures collected at hatch. We estimated a predicted growth curve only from our top model as the next best model had a ΔAIC_c_ > 6 (Appendix S1: Table S4). Chick growth did not differ among seasons, and our top-performing growth function included both a constant logistic coefficient (K = 0.13 ± 4.2×10^-3^) and inflection point (T_i_ *=* 17.5 ± 0.5 days).

The fit of our global model was greatest using 7-day averages of our continuous variables – daily biomass and daily median body size – and no random effects (Appendix S1: Table S5). Our top model explaining chick BCI (*n* = 89) included invertebrate biomass and hatch date with a smoothed age effect (*w_i_* = 0.75; Appendix S1: Table S6). Chick growth improved with higher invertebrate biomass (β = 1.8 × 10^-4^ ± 3.8 × 10^-5^ mg^-1^, CI = 1.2 × 10^-4^, 2.8 × 10^-4^; R^2^_adj._ = 0.37; Fig. 4a) but decreased with later hatch dates (β = -0.013 ± 0.003 d^-1^, CI = -0.0053, -0.019; Fig. 4b). Invertebrate body size had no consistent effect (Appendix S1: Fig. S3). Chicks had 3–75% higher body condition indices during periods with higher-than-average invertebrate biomass compared to periods with low invertebrate abundance. In terms of phenology, chicks grew better than expected if they hatched before 5 Jun but worse than expected if they hatched after that date.

**Figure 4.**
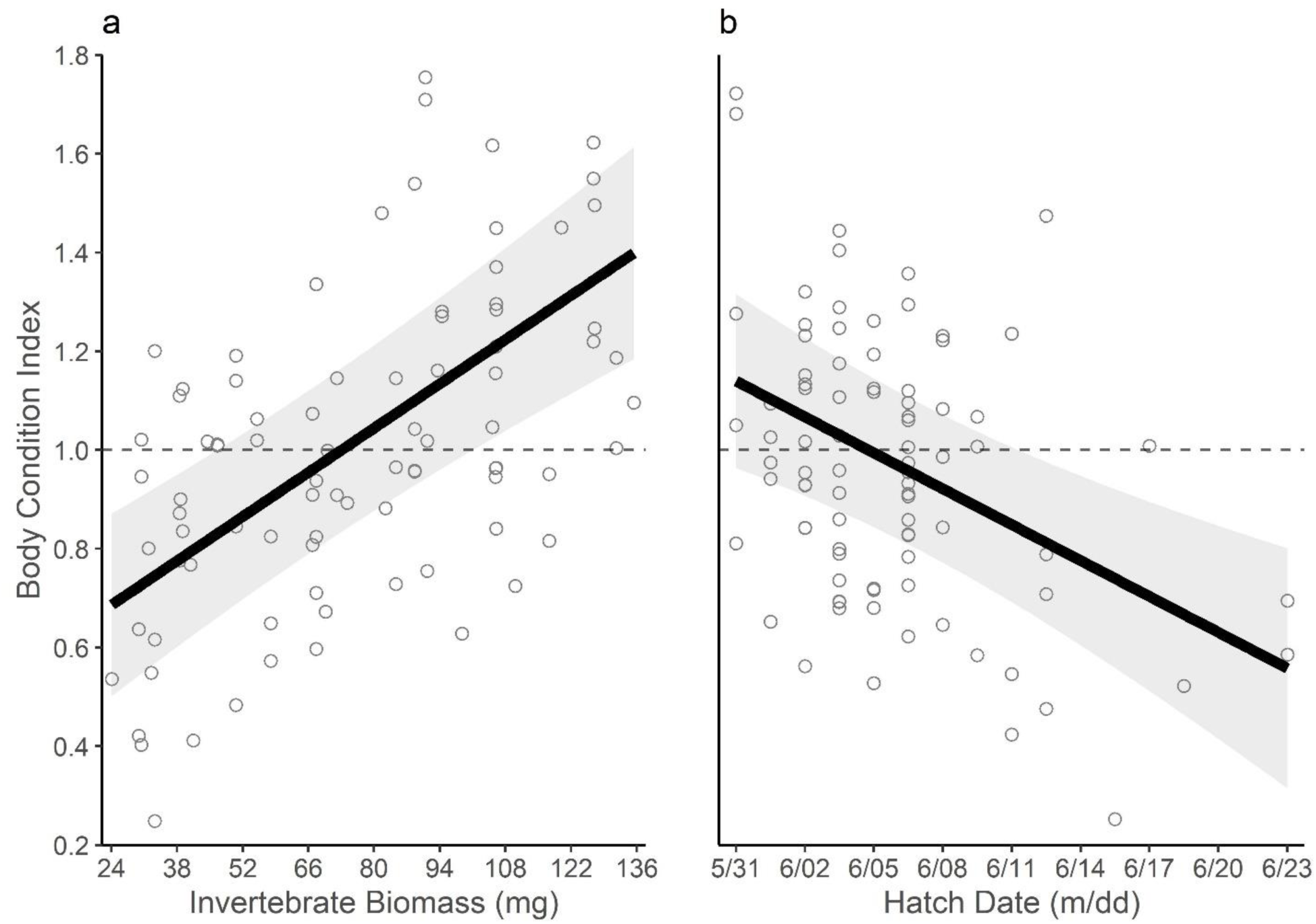
Effect of daily invertebrate biomass (a) and hatch date (b) on godwit chick body condition index (BCI). BCI (hollow points) is the ratio of the observed to expected weight gain since an individual’s last measurement. BCI > 1 correspond with above average growth and BCI < 1 below average growth (dashed line). Regression line (black) and 95% confidence interval (grey) are shown.

### Effect of resources on survival: constant or age-varying?

Of the 128 godwit chicks in our study, we excluded 6 due to human-caused mortality or instances when the radio fell off on the day of deployment. The mean DSR of the remaining 122 chicks was 86 ± 24%, meaning that 19.2 ± 33% survived to fledge, although this varied among seasons and broods (Appendix S1: Table S7).

The model with an age-varying effect of invertebrate body size (DIC = 282.2) outperformed the model with an age-varying effect of invertebrate biomass (DIC = 288.7). We therefore used the former in our subsequent models. The constant effect of invertebrate biomass and the age-varying invertebrate size effect had 79% and 85% posterior inclusion probabilities, respectively (Table 1). We also included constant effects of age and invertebrate size to accompany the interaction term.

**Table 1.** Bayesian model selection on variables in a global logistic model predicting daily survival rate in godwit chicks near Beluga River, AK from 2009–2019. Predictors were selected using the indicator-variable approach, in which posterior inclusion probabilities (weights) and Bayes Factors (BF) were estimated from a Bernoulli variable associated with each predictor. Variables of the global model with BF > 3 and their component parts (i.e., interaction terms) were included in the top model.

| Variable | Weight, global | BF, global | Weight, top | BF, top |
| --- | --- | --- | --- | --- |
| Age | 0.53 | 1.125 | 0.52 | 1.081 |
| Size | 0.54 | 1.177 | 0.52 | 1.068 |
| Hatch | 0.54 | 1.175 | - | - |
| Biomass | 0.79 | 3.730 | 0.77 | 3.340 |
| Age × Size | 0.85 | 5.614 | 0.80 | 4.042 |

*Age: number of days since chick hatched; Hatch: individual chick's hatch date; Size: daily median invertebrate body size; Biomass: daily invertebrate biomass*

Chick survival improved with greater invertebrate biomass and larger invertebrate body sizes, and the latter effect increased throughout development (Table 2). Each 1% increase in daily invertebrate biomass (+ 1.5 mg) improved daily chick survival by 0.66% (Fig. 5a), while each 1% increase in median invertebrate body size (+ 0.06 mg) led to a 1.02% increase in daily chick survival. This ‘size’ effect then grew by 2.2% with each day that a chick survived (Fig. 5b). Age itself, however, had no consistent effect on chick survival. Daily chick survival was, on average, 17% higher during periods of above average invertebrate biomass, and 29% higher in these instances for chicks below 5-days of age and 50–72% higher for chicks 11–21 d.

**Figure 5.**
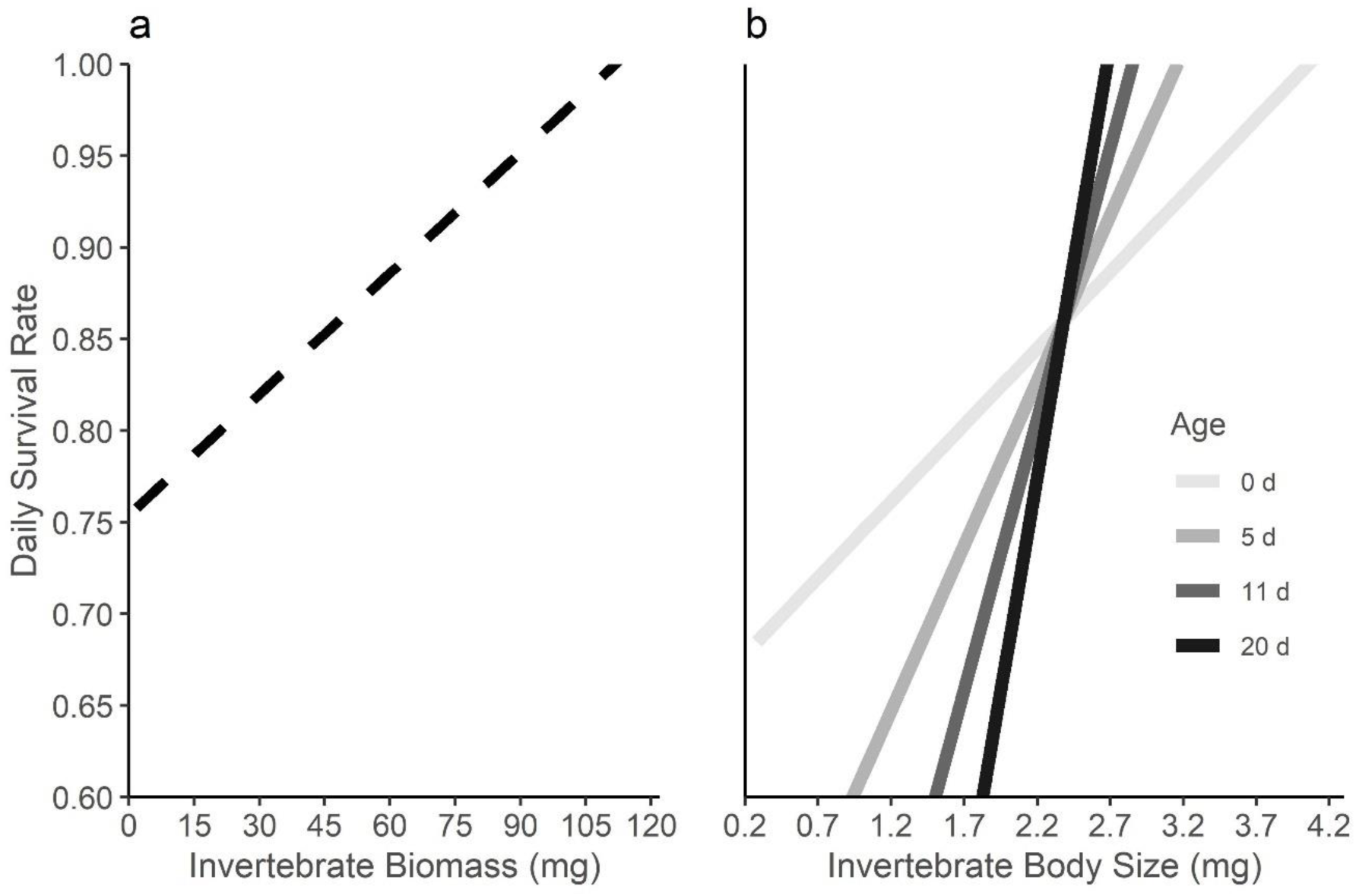
Effects of daily invertebrate biomass (a) and invertebrate body size (b) on the daily survival of godwit chicks from the posterior mean estimates of a Bayesian hierarchical model (credible intervals not shown). Biomass (dashed) had a constant effect, but the effect of size varied with age (shade of grey, in days).

**Table 2.**
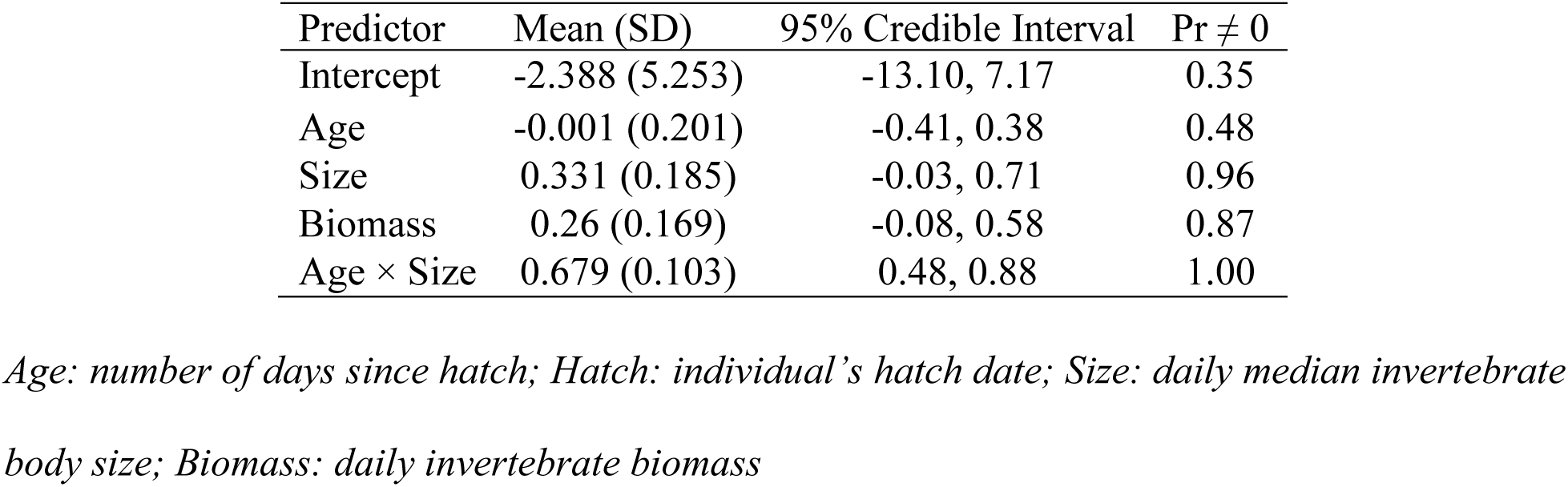
Standardized effect of variables on the survival rates of godwit chicks near Beluga River, AK from 2009–2019. Posterior probabilities were estimated from a hierarchical model (n = 122, posterior samples = 5000) with both survival and stochastic model components.

| Predictor | Mean (SD) | 95% Credible Interval | Pr ≠ 0 |
| --- | --- | --- | --- |
| Intercept | -2.388 (5.253) | -13.10, 7.17 | 0.35 |
| Age | -0.001 (0.201) | -0.41, 0.38 | 0.48 |
| Size | 0.331 (0.185) | -0.03, 0.71 | 0.96 |
| Biomass | 0.26 (0.169) | -0.08, 0.58 | 0.87 |
| Age × Size | 0.679 (0.103) | 0.48, 0.88 | 1.00 |

*Age: number of days since hatch; Hatch: individual’s hatch date; Size: daily median invertebrate body size; Biomass: daily invertebrate biomass*

### Population match and reproductive success

The model fit for the whole demand curve (AIC_c_ = -300.1) was 25.7-times better than the peak demand curve (AIC_c_ = –248.7). Godwits had, on average, 51.9 ± 9.2% overlap with resource phenology according to the peak demand model, but 44.7 ± 11.6% overlap according to the whole demand model. Seasons also differed in curve height (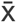 = 8,800 ± 3,668 mg) and the difference between the peak dates of the resource and demand curves (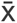 = 14.7 ± 16.36 d; Appendix S1: Fig. S4, S5).

Godwit fledging rates did not decrease linearly through time (Appendix S1: Table S8) but were lowest in 2014 and 2015. Those years had ∼19% poorer overlap and ∼28 days greater mismatching compared to the long-term average. Mismatching on this scale resulted in 24% lower fledging rates and near complete reproductive failure for the population. Models differed in their ability to explain population-level reproductive success but the whole demand model was best supported (Appendix S1: Table S9). The whole demand model explained 55% of the variation in godwit fledging rates (β = 1.19 ± 0.41; R^2^_adj._ = 0.55; *w_i_* = 0.43; Fig. 6 upper left; Appendix S1: Fig. S4). The ‘difference in peak dates’ model performed similarly well (β = -0.68 ± 0.27; R^2^_adj_ = 0.48; *w*_i_ = 0.36; Fig. 6 upper left) but was 7% less likely to be the top model. Both the peak demand overlap (β = 1.00 ± 0.56; R^2^_adj._ = 0.26; *w_i_* = 0.11; Fig. 6 lower left; Appendix S1: Fig. S5) and curve height models (β = 2.49 ± 1.44; R^2^_adj_ = 0.25; *w*_i_ = 0.10; Fig. 6 lower right) were unlikely to be the top model given their low model weights and the low amounts of interannual variation in fledging rates explained by either.

**Figure 6.**
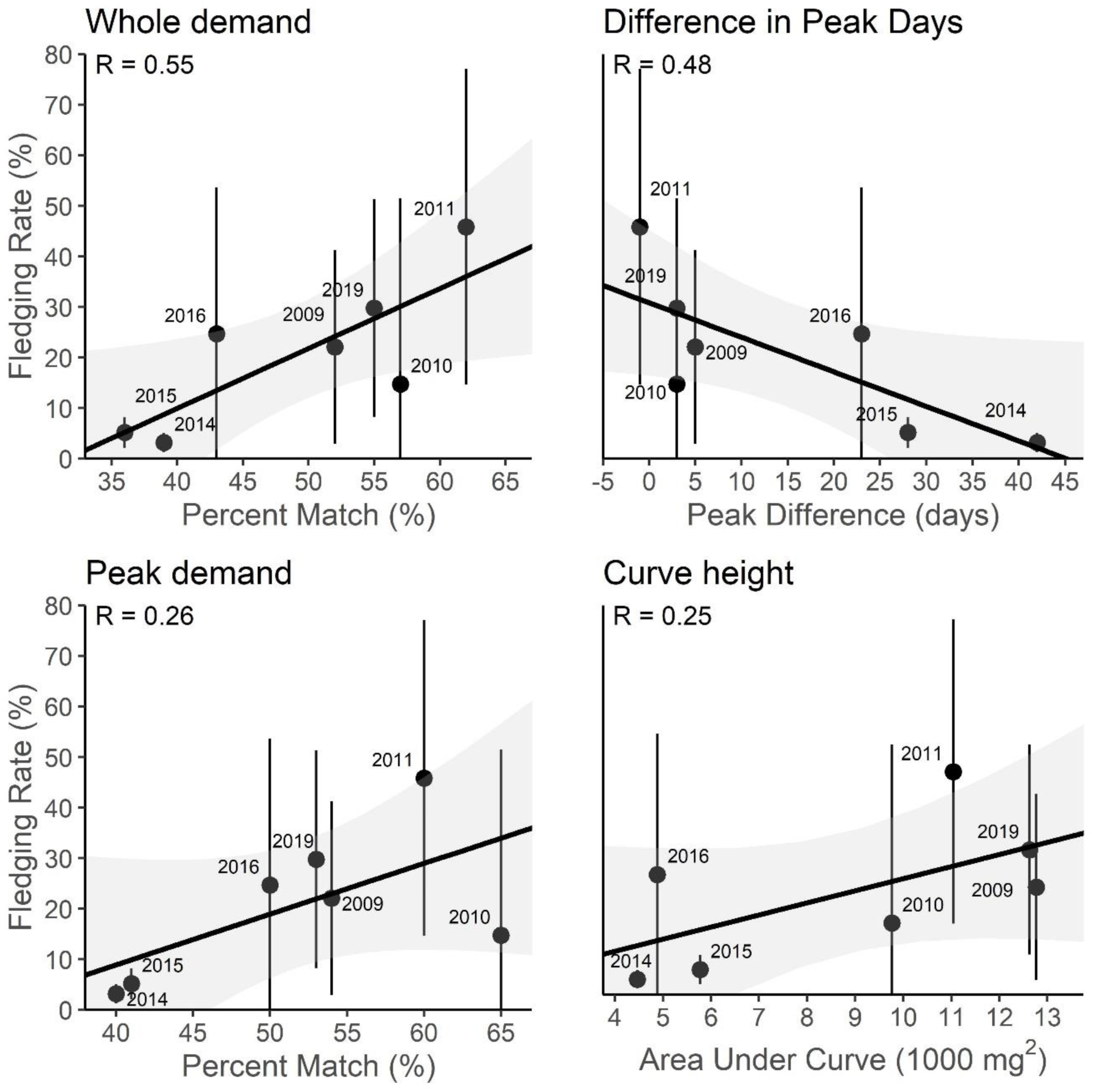
Correlation of seasonal fledging rates with measures of whole demand overlap (upper left), difference in peak dates (upper right), peak demand overlap (lower left), and curve height (lower right). Seasonal fledging rates were extrapolated from daily survival rates with accompanying 95% credible intervals. Regression lines (black) with 95% confidence interval (grey) are shown. Correlation coefficient for univariate models is displayed for each model.

## Discussion

The disconnect between empirical results and the theoretical predictions of the match-mismatch hypothesis make it difficult to assess the effects of climate change-induced phenological mismatches on consumer populations (Visser and Gienapp 2019, Keogan et al. 2020). To remedy this gap and connect mismatches to demographic processes, Kharouba & Wolkovich (2020) urged researchers to define pre-climate change baselines, collect per-capita data on resources and consumers, and test competing biological mechanisms. We developed mismatch models aimed at fulfilling these recommendations while adopting an age-structured representation of consumer demand. Using this approach, we built upon the findings of Senner et al. (2017) and identified heretofore undetected individual- and population-level fitness effects of mismatching in the Alaskan breeding population of Hudsonian godwits. Our study joins the growing literature suggesting that mismatches do not fall neatly into a ‘matched’ or ‘mismatched’ paradigm (Keogan et al. 2020, Simmonds et al. 2020). Instead, models built around the underlying biological mechanisms connecting consumers and resources are key to clarifying how mismatching affects consumer fitness (Takimoto and Sato 2020).

### More than mistiming: the tandem drivers of resource availability

We found that resources affected godwit chick survival in two distinct ways: first, periods with reduced resource abundance resulted in poorer growth and lower survival and, second, access to larger invertebrates was increasingly important to the survival of older chicks. Our findings differ from those of previous godwit studies, which found no effects of limited resource availability in the Alaskan godwit breeding population (Senner et al., 2017; Wilde et al., in revision). While these studies did not investigate the influence of invertebrate body size on godwit chicks, our contradictory conclusions likely stem from our use of hierarchical models that can approximate time-varying effects on survival (Royle and Dorazio 2009). Increasing energetic demands throughout ontogeny mean that the effects of resource limitation are unlikely to be constant over an individual’s lifetime (Yang and Rudolf 2010, Takimoto and Sato 2020). Therefore, models that accommodate varying predictor effects may be key to clarifying how resource characteristics affect consumer fitness.

Godwit chicks had improved growth and survival following periods with high resource abundance. Having adequate resources during energetically demanding periods is a primary driver of animal fitness (Bastille-Rousseau et al. 2015), especially in highly seasonal environments (McKinnon et al. 2012). Given their high energetic demands and rapid development, chicks of shorebird species across the Arctic exhibit survival costs following reduced resource abundance (Schekkerman et al. 2003, Saalfeld et al. 2019). Accordingly, godwit chicks in this study had higher body condition indices and higher daily survival probabilities during periods of higher-than-average invertebrate abundance. While we also detected effects of hatch date (i.e., phenology) on chick growth, these did not translate into an effect on survival. Our results therefore suggest that relating fitness measures to resource availability captures the effects of mismatching while defining its specific costs in biological terms (Dunn et al. 2011).

In addition to the effects of resource abundance, the quality (i.e., daily median body size) of invertebrates became increasingly important as godwit chicks aged. Optimal foraging theory predicts that consumers should select resources with the most energy content relative to foraging effort (Krebs et al. 1977). Chicks of black-tailed godwits (*Limosa limosa*), for instance, prioritize the rapid intake of small prey early in life, but switch to the slower intake of larger prey as they grow older (Schekkerman and Boele 2009). While we did not observe foraging behaviors directly, we hypothesize that Hudsonian godwit chicks may make a similar transition and increasing selection for larger prey could explain the especially high costs of poor resource quality for older chicks we found. Changes in resource quality, though rarely explored in the context of mismatches, can enact strong selection on consumer populations (Keogan et al. 2020, Yang et al. 2020). Because some individuals will encounter high-quality conditions in seasons when they are ‘mismatched’ (Kerby et al. 2012), accounting for the effects of multiple aspects of resource availability could improve our ability to document the true effects of mismatching.

Taken together, the additive effects of resource quantity and quality are likely to worsen in Beluga given the changes we observed in the invertebrate community. Climate-induced reductions in resource availability are common across terrestrial and marine systems (Bowden et al. 2015, Weterings et al. 2018). Arctic invertebrates, in particular, are simultaneously emerging earlier (Høye et al. 2007), becoming less abundant (van Klink et al. 2020), and smaller in size (Bowden et al. 2015, Jonsson et al. 2015) with increasing spring temperatures. Here, we found a linear decrease in the daily abundance and daily median body size of invertebrates, but no change in the date of peak occurrence of invertebrates over the course of our study. Although we did not detect a change in the timing of the resource peak, we found opposing trends over time in the abundance of invertebrates during the early and late portions of the godwit breeding season. Therefore, should these trends continue, developing godwit chicks may face increasingly untenable conditions as food becomes both less abundant and poorer in quality (i.e., smaller size). More broadly, our results suggest that resource timing, quality, and quantity can act as concomitant drivers of phenological mismatches (Rollins and Benard 2020), and that their effects may be most apparent when placed in the context of the consumer life cycle (Yang et al. 2020).

### Modelling the demand-resource interaction clarifies the population effects of mismatching

Variation in godwit reproductive success at the population level was best explained by our whole demand model of mismatches, although the simpler difference in dates model also performed well. Estimates from overlap and dates models often correlate (Ramakers et al. 2020), but may perform differently depending on a species’ life history and trophic specialization (Miller-Rushing et al. 2010). Thus, while difference in dates models may suffice for godwits and other species with narrow, synchronous breeding phenologies or those that rely on singular resource pulses (Miller-Rushing et al. 2010), they would likely perform poorly in species with highly variable nest initiation dates or those capable of multiple nesting events (Phillimore et al. 2016). Furthermore, difference in dates models could prove less accurate than whole demand models when resource phenology is multimodal or lacks a clearly defined peak, such as those typical of the temperate zone (Pearce-Higgins et al. 2005). Because overlap models account for both synchrony and the magnitude of available energy between interacting consumer-resource pairs, they are more likely to capture mismatching as a disrupted interaction (Kerby et al. 2012). Overlap models are therefore likely more generalizable, but using both overlap and difference in dates models could help when exploring how mismatching occurs on a case-by-case basis (Kellermann and van Riper 2015).

Not all overlap models are equivalent, however. Overlap models have received mixed support (Ramakers et al. 2020), but their ability to accurately quantify mismatching at the tails of the consumer curve has been suggested as an important component of their effectiveness (Kerby et al. 2012). Whereas our peak demand model performed relatively poorly, our whole demand model explained the most variation in fledging rates among our suite of models. The difference between the two models’ performance likely stems from the inability of the peak demand model to accurately capture consumer demand at the upper (i.e., right-hand) tail of the consumer curve, corresponding to the period when individual-level consumer demand is greatest. Our results therefore show that incorporating additional nuance into the statistical concept of consumer phenologies can greatly improve overlap models (Lindén 2018).

The need to accurately identify mismatches is made most clear by the accumulating evidence for variable and non-linear responses by consumer populations to mismatching (Visser and Both 2005, Phillimore et al. 2016). So called ‘tipping points’ – thresholds past which an effect abruptly changes (Latty and Dakos 2019) – buffer consumer populations from the negative impacts of moderate mismatching and may contribute to the lack of consistent responses to mismatching across consumer populations (Simmonds et al. 2020). In this population of godwits, we found that greater population-level mismatching consistently drove poorer fledging success, but that there may be thresholds past which the effects are most severe. For instance, the degree of mismatch in 2014 and 2015 resulted in near complete reproductive failure for the population. Similarly low fledging rates for Hudson Bay breeding godwits, which are mismatched by 11- days on-average (Senner et al. 2017), suggests that for godwits, this tipping point may exist when populations are mismatched by more than ∼10 days or have less than 40% overlap with the resource curve.

Importantly though, the 2014 and 2015 seasons in Beluga coincided with a period of anomalous and prolonged near-surface warming in the northeastern Pacific called the ‘blob’ (Cavole et al. 2016). Thus, while the conditions in these atypical seasons may provide useful insights into potential outcomes of a warming climate on coastal communities in the region (Auth et al. 2018), mismatches of this magnitude are unlikely to become the norm. Beluga godwits have been able to advance their timing of migration and reproduction in response to recent long-term, linear warming trends (Senner, 2012; Senner et al., 2017). Their ability to do so into the future will depend on whether the cues godwits use to time their annual cycle remain predictive of the timing of resource pulses on their breeding grounds. The significant spring warming and earlier snow disappearance dates projected for the North American sub-Arctic, for instance, means that godwits and other migratory populations may soon face accelerating, potentially non-linear warming to which they have limited capacity to respond (Love et al. 2010, Lader et al. 2020). Quantifying the strength and effects of mismatching in real time will thus be crucial for conservation going forward (Simmonds et al. 2020).

### Conclusions

By modelling the unspoken assumptions of the match-mismatch hypothesis, we stand to adopt a more powerful definition of mismatching in biological terms and, in so doing, be better able to identify the circumstances under which consumer populations perform poorly. Our work also illustrates the role of ontogeny in shaping an individual’s changing response to resource availability over time (Yang and Rudolf 2010), and helps explain the empirical-theoretical disconnect in phenological studies. Importantly, our models are transferrable to other systems, whereby remotely-sensed indices and knowledge of a population’s age-structure could approximate resource availability and energetic requirements, respectively, when these data are otherwise unavailable (Lumbierres et al. 2017). Finally, we show how treating mismatches as an outcome of both consumer demand and resource dynamics provides insight into the structure of individual-level effects and the mechanism behind population-level responses (Takimoto and Sato 2020). Replacing the categorical ‘matched’ or ‘mismatched’ view of mismatching with one that explicitly recognizes the underlying mechanism may be critical to monitoring and conserving animal populations in an uncertain future.

## Acknowledgements

We thank the Cook Inlet Region Inc. for permitting us access to their lands. We thank Drs. Sarah Converse and Emily Weiser for analytical guidance, and Drs. Sarah Saalfield and Jay VonBank for comments on early drafts. Funding was provided by the Association of Field Ornithologists, Wilson Ornithological Society, American Ornithological Society, Arctic Audubon Society, Cornell Lab of Ornithology, Athena Fund at the Cornell Lab of Ornithology, University of South Carolina, Faucett Family Foundation, the David and Lucile Packard Foundation, National Science Foundation (PCE-1110444 and DGE-1144153), and the U.S. Fish and Wildlife Service (4074, 5147). All procedures met the ethical standards of the University of South Carolina (2449-101417-042219), Alaska Department of Fish and Game (20-024), and USGS (24191). Any use of trade, product, or firm names is for descriptive purposes only and does not imply endorsement by the U.S. Government. The authors declare no conflict of interest.

## Author Contributions

LRW and NRS conceived of the study. All authors collected field data. LRW analyzed the data and wrote the manuscript. All authors contributed to revisions.

## Data Availability Statement

Data are deposited at Dryad (https://doi.org/10.5061/dryad.x69p8czh0) and computer code for all analyses are available at github (http://doi.org/10.5281/zenodo.4298755).

## References

Auth, T. D., E. A. Daly, R. D. Brodeur, and J. L. Fisher. 2018. Phenological and distributional shifts in ichthyoplankton associated with recent warming in the northeast Pacific Ocean. Global Change Biology 24:259–272.

Bartoń, K. 2015. Multiple-Model Inference (Package ‘MuMIn’). R v. 3.3.2.

Bastille-Rousseau, G., J. R. Potts, J. A. Schaefer, M. A. Lewis, E. H. Ellington, N. D. Rayl, S. P. Mahoney, and D. L. Murray. 2015. Unveiling trade-offs in resource selection of migratory caribou using a mechanistic movement model of availability. Ecography 38:1049–1059.

Bates D. M. Maechler B. Bolker and S. Walker. 2015. Fitting linear mixed-effects models using lme4. Journal of Statistical Software, 67: 1–48.

Both, C., M. V. Asch, R. G. Bijlsma, A. B. V. D. Burg, and M. E. Visser. 2009. Climate change and unequal phenological changes across four trophic levels: constraints or adaptations? Journal of Animal Ecology 78:73–83.

Both, C., and M. E. Visser. 2001. Adjustment to climate change is constrained by arrival date in a long-distance migrant bird. Nature 411:296–298.

Bowden, J. J., A. Eskildsen, R. R. Hansen, K. Olsen, C. M. Kurle, and T. T. Høye. 2015. High-Arctic butterflies become smaller with rising temperatures. Biology Letters 11:20150574.

Brown, S. C., H. R. Gates, J. R. Liebezeit, P. A. Smith, B. L. Hill, and R. B. Lanctot. 2014. Arctic Shorebird Demographics Network Breeding Camp Protocol, Version 5. Unpubl. paper by U.S. Fish and Wildlife Service and Manomet Center for Conservation Sciences:118.

Burnham, K. P., and D. R. Anderson. 2002. Model selection and multimodel inference: A practical information–theoretic approach. Springer Science and Business Media, New York.

Cavole, L. M., A. M. Demko, R. E. Diner, A. Giddings, I. Koester, C. M. L. S. Pagniello, M.-L. Paulsen, A. Ramirez-Valdez, S. M. Schwenck, N. K. Yen, M. E. Zill, and P. J. S. Franks. 2016. Biological Impacts of the 2013–2015 Warm-Water Anomaly in the Northeast Pacific: Winners, Losers, and the Future. Oceanography 29:273–285.

Chmura, H. E., H. M. Kharouba, J. Ashander, S. M. Ehlman, E. B. Rivest, and L. H. Yang. 2019. The mechanisms of phenology: the patterns and processes of phenological shifts. Ecological Monographs 89:e01337.

Converse, S. J., J. A. Royle, P. H. Adler, R. P. Urbanek, and J. A. Barzen. 2013. A hierarchical nest survival model integrating incomplete temporally varying covariates. Ecology and Evolution 3:4439–4447.

Corkery, C. A., E. Nol, and L. Mckinnon. 2019. No effects of asynchrony between hatching and peak food availability on chick growth in Semipalmated Plovers (Charadrius semipalmatus) near Churchill, Manitoba. Polar Biology 42:593–601.

Cushing, D. H. 1974. The natural regulation of fish populations. Pages 399–412 Sea Fisheries Research. F. R. Harden Jones, ed, London: Elek Science.

Cushing, D. H. 1990. Plankton Production and Year-class Strength in Fish Populations: an Update of the Match/Mismatch Hypothesis. Pages 249–293 in J. H. S. Blaxter and A. J. Southward, editors. Advances in Marine Biology. Academic Press.

Denwood, M. J. 2016. runjags: An R Package Providing Interface Utilities, Model Templates, Parallel Computing Methods and Additional Distributions for MCMC Models in JAGS. Journal of Statistical Software, 71: 1–25.

Dunn, P. O., D. W. Winkler, L. A. Whittingham, S. J. Hannon, and R. J. Robertson. 2011. A test of the mismatch hypothesis: How is timing of reproduction related to food abundance in an aerial insectivore? Ecology 92:450–461.

Ganihar, S. R. 1997. Biomass estimates of terrestrial arthropods based on body length. Journal of Biosciences 22:219–224.

Gelman, A. 2008. Scaling regression inputs by dividing by two standard deviations. Statistics in Medicine 27:2865–2873.

Gelman, A., and D. B. Rubin. 1992. Inference from Iterative Simulation Using Multiple Sequences. Statistical Science 7:457–472.

Høye, T. T., E. Post, H. Meltofte, N. M. Schmidt, and M. C. Forchhammer. 2007. Rapid advancement of spring in the High Arctic. Current Biology 17:R449–R451.

Jonsson, M., P. Hedström, K. Stenroth, E. R. Hotchkiss, F. R. Vasconcelos, J. Karlsson, and P. Byström. 2015. Climate change modifies the size structure of assemblages of emerging aquatic insects. Freshwater Biology 60:78–88.

Kellermann, J. L., and C. van Riper. 2015. Detecting mismatches of bird migration stopover and tree phenology in response to changing climate. Oecologia 178:1227–1238.

Keogan, K., S. Lewis, R. J. Howells, M. A. Newell, M. P. Harris, S. Burthe, R. A. Phillips, S. Wanless, A. B. Phillimore, and F. Daunt. 2020. No evidence for fitness signatures consistent with increasing trophic mismatch over 30 years in a population of European shag Phalacrocorax aristotelis. Journal of Animal Ecology n/a.

Kerby, J. T., C. C. Wilmers, and E. Post. 2012. Climate change, phenology and the nature of consumer–resource interactions: advancing the match/mismatch hypothesis. Pages 508– 525 Trait-mediated indirect interactions: ecological and evolutionary perspectives (eds Ohgushi T, Schmitz O, Holt R). Cambridge University Press, Cambridge.

Kharouba, H. M., and E. M. Wolkovich. 2020. Disconnects between ecological theory and data in phenological mismatch research. Nature Climate Change 10:406–415.

van Klink, R., D. E. Bowler, K. B. Gongalsky, A. B. Swengel, A. Gentile, and J. M. Chase. 2020. Meta-analysis reveals declines in terrestrial but increases in freshwater insect abundances. Science 368:417–420.

Krebs, C. J., J. T. Erichson, M. I. Weber, and E. I. Charnov. 1977. Optimal prey selection in the great tit (Parus major). Animal Behaviour 25:30–38.

Kwon, E., E. L. Weiser, R. B. Lanctot, S. C. Brown, H. R. Gates, G. Gilchrist, S. J. Kendall, D. B. Lank, J. R. Liebezeit, L. McKinnon, E. Nol, D. C. Payer, J. Rausch, D. J. Rinella, S. T. Saalfeld, N. R. Senner, P. A. Smith, D. Ward, R. W. Wisseman, and B. K. Sandercock. 2019. Geographic variation in the intensity of warming and phenological mismatch between Arctic shorebirds and invertebrates. Ecological Monographs 89:e01383.

Lader, R., J. E. Walsh, U. S. Bhatt, and P. A. Bieniek. 2020. Anticipated changes to the snow season in Alaska: Elevation dependency, timing and extremes. International Journal of Climatology 40(1):169–187. Journal of Climatology 40:169–187.

Lameris, T. K., I. Scholten, S. Bauer, M. M. P. Cobben, B. J. Ens, and B. A. Nolet. 2017. Potential for an Arctic-breeding migratory bird to adjust spring migration phenology to Arctic amplification. Global Change Biology 23:4058–4067.

Latty, T., and V. Dakos. 2019. The risk of threshold responses, tipping points, and cascading failures in pollination systems. Biodiversity and Conservation 28:3389–3406.

Leung, M. C.-Y., E. Bolduc, F. I. Doyle, D. G. Reid, B. S. Gilbert, A. J. Kenney, C. J. Krebs, and J. Bêty. 2018. Phenology of hatching and food in low Arctic passerines and shorebirds: is there a mismatch? Arctic Science 4:538–556.

Liebezeit, J. R., P. A. Smith, R. B. Lanctot, H. Schekkerman, I. Tulp, S. J. Kendall, D. M. Tracy, R. J. Rodrigues, H. Meltofte, J. A. Robinson, C. Gratto-Trevor, B. J. Mccaffery, J. Morse, and S. W. Zack. 2007. Assessing the Development of Shorebird Eggs Using the Flotation Method: Species-Specific and Generalized Regression Models. The Condor 109:32–47.

Lindén, A. 2018. Adaptive and nonadaptive changes in phenological synchrony. Proceedings of the National Academy of Sciences 115:5057–5059.

Link, W. A., and R. J. Barker. 2006. Model Weights and the Foundations of Multimodel Inference. Ecology 87:2626–2635.

Loonstra, A. H. J., M. A. Verhoeven, and T. Piersma. 2018. Sex-specific growth in chicks of the sexually dimorphic Black-tailed Godwit. Ibis 160:89–100.

Love, O. P., H. G. Gilchrist, S. Descamps, C. A. D. Semeniuk, and J. Bêty. 2010. Pre-laying climatic cues can time reproduction to optimally match offspring hatching and ice conditions in an Arctic marine bird. Oecologia 164:277–286.

Lumbierres, M., P. Méndez, J. Bustamante, R. Soriguer, and L. Santamaría. 2017. Modeling Biomass Production in Seasonal Wetlands Using MODIS NDVI Land Surface Phenology. Remote Sensing 9:392.

Maechler, M. 2020. sfsmisc: Utilities from ’Seminar fuer Statistik’ ETH Zurich. Version 1.1–7.

McKinnon, L., M. Picotin, E. Bolduc, C. Juliet, and J. Bety. 2012. Timing of breeding, peak food availability, and effects of mismatch on chick growth in birds nesting in the High Arctic. Canadian Journal of Zoology 90:961–971.

Miller-Rushing, A. J., T. T. Høye, D. W. Inouye, and E. Post. 2010. The effects of phenological mismatches on demography. Philosophical Transactions of the Royal Society B: Biological Sciences 365:3177–3186.

Parmesan, C., and G. Yohe. 2003. A globally coherent fingerprint of climate change impacts across natural systems. Nature 421:37–42.

Pearce-Higgins, J. W., D. W. Yalden, and M. J. Whittingham. 2005. Warmer Springs Advance the Breeding Phenology of Golden Plovers Pluvialis apricaria and Their Prey (Tipulidae). Oecologia 143:470–476.

Phillimore, A. B., D. I. Leech, J. W. Pearce-Higgins, and J. D. Hadfield. 2016. Passerines may be sufficiently plastic to track temperature-mediated shifts in optimum lay date. Global Change Biology 22:3259–3272.

Pinheiro, J., and D. Bates. 2000. Mixed-Effects Models in S and S-PLUS. Springer-Verlag, New York.

Pinheiro, J., D. Bates, S. DebRoy, D. Sarkar, and R Core Team. 2020. nlme: Linear and Nonlinear Mixed Effects Models. R package 3:1–149.

Plummer, M. 2012. JAGS: just another Gibbs sampler. Version 3.3.0.

Plummer, M. 2013. Package rjags: Bayesian graphical models using MCMC. Version 3.10.

Powell, L. A. 2007. Approximating variance of demographic parameters using the delta method: a reference for avian biologists. The Condor 109:949–954.

R Core Team (2020) R: A Language and Environment for Statistical Computing. R Foundation for Statistical Computing, Vienna, Austria.

Ramakers, J. J. C., P. Gienapp, and M. E. Visser. 2020. Comparing two measures of phenological synchrony in a predator–prey interaction: Simpler works better. Journal of Animal Ecology 89:745–756.

Reed, T. E., S. Jenouvrier, and M. E. Visser. 2013. Phenological mismatch strongly affects individual fitness but not population demography in a woodland passerine. Journal of Animal Ecology 82:131–144.

Rigby, R. A., and D. M. Stasinopoulos. 2005. Generalized additive models for location, scale and shape. Journal of the Royal Statistical Society 54:507–554.

Rogers, L. E., R. L. Buschbom, and C. R. Watson. 1977. Length-Weight Relationships of Shrub-Steppe Invertebrates1. Annals of the Entomological Society of America 70:51–53.

Rollins, H. B., and M. F. Benard. 2020. Challenges in predicting the outcome of competition based on climate change-induced phenological and body size shifts. Oecologia 193:749– 759.

Royle, J. A., and R. M. Dorazio. 2009. Hierarchical Modeling and Inference in Ecology, The Analysis of Data from Populations, Metapopulations and Communities.

Saalfeld, S. T., D. C. McEwen, D. C. Kesler, M. G. Butler, J. A. Cunningham, A. C. Doll, W. B. English, D. E. Gerik, K. Grond, P. Herzog, B. L. Hill, B. J. Lagassé, and R. B. Lanctot. 2019. Phenological mismatch in Arctic-breeding shorebirds: Impact of snowmelt and unpredictable weather conditions on food availability and chick growth. Ecology and Evolution 9:6693–6707.

Samplonius, J. M., L. Bartošová, M. D. Burgess, A. V. Bushuev, T. Eeva, E. V. Ivankina, A. B. Kerimov, I. Krams, T. Laaksonen, M. Mägi, R. Mänd, J. Potti, J. Török, M. Trnka, M. E. Visser, H. Zang, and C. Both. 2018. Phenological sensitivity to climate change is higher in resident than in migrant bird populations among European cavity breeders. Global Change Biology 24:3780–3790.

Schekkerman, H., and A. Boele. 2009. Foraging in precocial chicks of the black-tailed godwit Limosa limosa: vulnerability to weather and prey size. Journal of Avian Biology 40:369– 379.

Schekkerman, H., I. Tulp, T. Piersma, and G. H. Visser. 2003. Mechanisms promoting higher growth rate in arctic than in temperate shorebirds. Oecologia 134:332–342.

Schekkerman, H., and G. H. Visser. 2001. Prefledging Energy Requirements in Shorebirds: Energetic Implications of Self-Feeding Precocial Development. The Auk 118:944–957.

Senner, N. R. 2012. One species but two patterns: Populations of the Hudsonian Godwit (*Limosa haemastica* ) differ in spring migration timing. The Auk 129:670–682.

Senner, N. R., M. Stager, and B. K. Sandercock. 2017. Ecological mismatches are moderated by local conditions for two populations of a long-distance migratory bird. Oikos 126:61–72.

Sharpe, F., M. Bolton, R. Sheldon, and N. Ratcliffe. 2009. Effects of Color Banding, Radio Tagging, and Repeated Handling on the Condition and Survival of Lapwing Chicks and Consequences for Estimates of Breeding Productivity. Journal of Field Ornithology 80:101–110.

Simmonds, E. G., E. F. Cole, B. C. Sheldon, and T. Coulson. 2020. Phenological asynchrony: a ticking time-bomb for seemingly stable populations? Ecology Letters.

Swift, R. J., A. D. Rodewald, and N. R. Senner. 2017a. Environmental heterogeneity and biotic interactions as potential drivers of spatial patterning of shorebird nests. Landscape Ecology 32:1689–1703.

Swift, R. J., A. D. Rodewald, and N. R. Senner. 2017b. Breeding habitat of a declining shorebird in a changing environment. Polar Biology 40:1777–1786.

Swift, R. J., A. D. Rodewald, and N. R. Senner. 2018. Context-dependent costs and benefits of a heterospecific nesting association. Behavioral Ecology 29:974–983.

Takimoto, G., and T. Sato. 2020. Timing and duration of phenological resources: Toward a mechanistic understanding of their impacts on community structure and ecosystem processes in stream food chains. Ecological Research 35:463–473.

Tulp, I., and H. Schekkerman. 2008. Has Prey Availability for Arctic Birds Advanced with Climate Change? Hindcasting the Abundance of Tundra Arthropods Using Weather and Seasonal Variation 61:48–60.

Vatka, E., M. Orell, and S. Rytkönen. 2016. The relevance of food peak architecture in trophic interactions. Global Change Biology 22:1585–1594.

Visser, M. E., and C. Both. 2005. Shifts in phenology due to global climate change: the need for a yardstick. Proceedings of the Royal Society B: Biological Sciences 272:2561–2569.

Visser, M. E., and P. Gienapp. 2019. Evolutionary and demographic consequences of phenological mismatches. Nature Ecology & Evolution 3:879–885.

Visser, M. E., L. te Marvelde, and M. E. Lof. 2012. Adaptive phenological mismatches of birds and their food in a warming world. Journal of Ornithology 153:75–84.

Visser, M. E., A. J. van Noordwijk, J. M. Tinbergen, and C. M. Lessells. 1998. Warmer springs lead to mistimed reproduction in great tits (Parus major). Proceedings of the Royal Society of London. Series B: Biological Sciences 265:1867–1870.

Walker, B. M., N. R. Senner, C. S. Elphick, and J. Klima. 2011. Hudsonian Godwit (Limosa haemastica), version 2.0. Page *in* A. F. Poole, editor. The Birds of North America.

Weterings, M. J. A., S. Moonen, H. H. T. Prins, S. E. van Wieren, and F. van Langevelde. 2018. Food quality and quantity are more important in explaining foraging of an intermediate-sized mammalian herbivore than predation risk or competition. Ecology and Evolution 8:8419–8432.

Wilde, L. R., R. J. Swift, and N. R. Senner. in review. Flexible space use and density-dependent heterospecific interactions determine fledging success in a precocial bird. Journal of Animal Ecology.

Williams, J. B., B. I. Tieleman, G. H. Visser, and R. E. Ricklefs. 2007. Does Growth Rate Determine the Rate of Metabolism in Shorebird Chicks Living in the Arctic? Physiological and Biochemical Zoology 80:500–513.

Yang, L. H., M. L. Cenzer, L. J. Morgan, and G. W. Hall. 2020. Species-specific, age-varying plant traits affect herbivore growth and survival. Ecology 101:e03029.

Yang, L. H., and V. H. W. Rudolf. 2010. Phenology, ontogeny and the effects of climate change on the timing of species interactions. Ecology Letters 13:1–10.

