## Appendix S1 for "The anatomy of a phenological mismatch: interacting consumer demand and resource characteristics determine the consequences of mismatching"

Table S1. Estimates of Pearson’s correlation coefficient (r) among pairs of fixed effect covariates in a generalized additive mixed model (GAMM) predicting godwit chick body condition index (BCI) estimates collected in Beluga River, Alaska from 2009–2019.

| Variable 1 | Variable 2 | r | lower | upper |
| --- | --- | --- | --- | --- |
| Size | Insect | 0.22 | 0.01 | 0.41 |
| Size | Hatch | 0.27 | 0.07 | 0.46 |
| Insect | Hatch | 0.04 | -0.17 | 0.24 |

Table S2. Estimates of Pearson’s correlation coefficient (r) among pairs of fixed effect covariates in a Bayesian hierarchical model on the survival of Hudsonian godwit chicks living near Beluga River, Alaska (2009 – 2019). Coefficients were measured to check collinearity between predictors in a global model prior to model selection procedures.

| Variable 1 | Variable 2 | r | lower | upper |
| --- | --- | --- | --- | --- |
| Size | Age | -0.006 | -0.04 | 0.03 |
| Size | Insect | 0.32 | 0.78 | 0.36 |
| Size | Hatch | -0.05 | -0.09 | -0.01 |
| Insect | Age | -0.03 | -0.08 | 0.02 |
| Insect | Hatch | 0.07 | 0.02 | 0.11 |
| Hatch | Age | -0.04 | -0.06 | -0.01 |

Table S3. Summary of season lengths and the observed Hudsonian godwit hatching dates and model predicted invertebrate resource peak from 2009 – 2019 near Beluga River, Alaska. The resource peak was calculated as the day with the smallest first derivative along the day + day^2^ curve.

| Year | Chicks  hatched | Season  length | Mean hatch  date (± SD) | Predicted resource  peak |
| --- | --- | --- | --- | --- |
| 2009 | 69 | 3 May – 10 Jul | 5 Jun (± 5.5 d) | 5 Jun |
| 2010 | 60 | 3 May – 10 Jul | 11 Jun (± 9.4 d) | 8 Jun |
| 2011 | 87 | 3 May – 10 Jul | 7 Jun (± 5.0 d) | 14 Jun |
| 2012^1^ | 32 | 8 May – 5 Jun | 9 Jun (± 5.0 d) | - |
| 2014 | 31 | 9 May – 13 Jul | 9 Jun (± 5.6 d) | 10 May |
| 2015 | 62 | 3 May – 10 Jul | 6 Jun (± 3.2 d) | 31 May |
| 2016 | 38 | 1 May – 10 Jul | 5 Jun (± 4.0 d) | 20 May |
| 2017^1^ | - | 11 May – 19 May | - | - |
| 2019 | 56 | 6 May – 26 Jul | 7 Jun (± 5.3 d) | 11 Jun |

^1^ Neither godwits or invertebrates were monitored for the full 2012 or 2017 season.

Table S4. Model selection table of logistic models predicting godwit chick mass from weekly captures (*n* = 103) of godwit chicks near Beluga River, Alaska from 2009 ̶ 2019. Following Senner et al. (2017) asymptotic mass was set to the adult average (~ 249 g). Initial values were set prior to modelling: inflection point T_i_ = 10.7, logistic coefficient K = 0.12. Chick identity (ID) was included as a random intercept for variables (✓), and all models had at least one random intercept term. Parameter estimates were averaged from 100 iterations. AIC_c_ = Akaike’s Information Criterion corrected for small sample sizes.

| Model no. | Variable | Value | Random  intercept | ΔAIC_c_ | log-likelihood | Model  weight (*w_i_*) | No. parameters |
| --- | --- | --- | --- | --- | --- | --- | --- |
| 2 | T_i_ | ~ 1 ` | ✓ | 0.0 | -528.0 | 0.98 | 6 |
|  | K | ~ 1 ` | ✓ |  |  |  |  |
| 4 | T_i_ | ~ year | ✓ | 6.9 | -528.5 | 0.02 | 9 |
|  | K | ~ 1 ` |  |  |  |  |  |
| 10 | T_i_ | ~ year | ✓ | 11.7 | -525.9 | >0.01 | 14 |
|  | K | ~ year |  |  |  |  |  |
| 1 | T_i_ | ~ 1 ` | ✓ | 17.5 | -538.3 | >0.01 | 4 |
|  | K | ~ 1 ` |  |  |  |  |  |
| 8 | T_i_ | ~ 1 ` | ✓ | 21.2 | -535.6 | >0.01 | 9 |
|  | K | ~ year |  |  |  |  |  |
| 5 | T_i_ | ~ year |  | 35.5 | -542.8 | >0.01 | 9 |
|  | K | ~ 1 ` | ✓ |  |  |  |  |
| 11 | T_i_ | ~ year |  | 40.0 | -540.1 | >0.01 | 14 |
|  | K | ~ year | ✓ |  |  |  |  |
| 7 | T_i_ | ~ 1 ` |  | 78.8 | -564.5 | >0.01 | 9 |
|  | K | ~ year | ✓ |  |  |  |  |
| 3 | T_i_ | ~ 1 ` |  | 80.7 | -570.4 | >0.01 | 4 |
|  | K | ~ 1 ` | ✓ |  |  |  |  |
| 9 | T_i_ | ~ 1 ` | ✓ | 136.4 | -593.1 | >0.01 | 9 |
|  | K | ~ year | ✓ |  |  |  |  |
| 12 | T_i_ | ~ year | ✓ | 183.5 | -602.4 | >0.01 | 14 |
|  | K | ~ year | ✓ |  |  |  |  |
| 6 | T_i_ | ~ year | ✓ | 188.4 | -605.2 | >0.01 | 9 |
|  | K | ~ 1 ` | ✓ |  |  |  |  |

Table S5. Model selection table of random intercept terms (top) and the timescale of continuous covariate (bottom) in our global generalized additive mixed model (GAMM) to predict body condition index (BCI) of godwit chicks near Beluga River, Alaska from 2009 ̶ 2019. Timescale is the period over which the continuous fixed effect variables – daily invertebrate biomass and daily median invertebrate body size – in the global model were averaged for model smoothing. AIC_c_ = Akaike’s Information Criterion corrected for small sample sizes.

| Random intercepts | | |  |  |
| --- | --- | --- | --- | --- |
|  | ΔAIC_c_ | Deviance | Model weight (*w_i_*) | No. parameters |
| ~ 1 | 0 | 11.6 | 0.42 | 4 |
| ~ 1\|chick | 4.2 | 14.5 | 0.22 | 59 |
| ~ 1\|brood + 1\|chick | 5.0 | 16.1 | 0.17 | 51 |
| ~ 1\|year | 5.2 | 17.5 | 0.10 | 9 |
| ~ 1\|year + 1\|brood | 7.8 | 19.0 | 0.04 | 52 |
| ~ 1\|brood | 8.1 | 19.3 | 0.03 | 47 |
| ~ 1\|year + 1\|chick/brood | 11.9 | 23.5 | >.01 | 64 |
| Timescale | | |  |  |
|  | ΔAIC_c_ | Deviance | Model weight (*w_i_*) | log-  Likelihood |
| 7-day avg. | 0 | 12.7 | 0.69 | -39.6 |
| 3-day avg. | 2.3 | 13.0 | 0.22 | -40.8 |
| day of | 4.7 | 134 | 0.06 | -42.0 |
| 1-day avg. | 6.4 | 13.6 | 0.03 | -42.8 |

Table S6. Comparison by AIC_c_ value among candidate models with predictor variables of Hudsonian godwit chick growth (*n* = 89) from 2009 – 2019, excluding chicks from 2014 (which lacked recaptures). Growth was estimated from body condition index (BCI) scores obtained from weekly captures. Continuous predictors were averaged over a 7-day period prior to BCI estimation. Inclusion in a model is indicated by a beta coefficient for predictors and plus signs (+) for smoothing terms. AIC_c_ = Akaike’s Information Criterion corrected for small sample sizes.

| Intercept | Invertebrate  biomass | Invertebrate  body size | Hatch date |  | s(Chick age) | df | Log-likelihood | AIC_c_ | ΔAIC_c_ | model weight |
| --- | --- | --- | --- | --- | --- | --- | --- | --- | --- | --- |
| 0.445 | 0.006 |  | -0.254 |  | + | 10 | -22.915 | 70.651 | 0 | 0.75 |
| 0.396 | 0.006 | 0.019 | -0.264 |  | + | 12 | -22.506 | 73.095 | 2.444 | 0.22 |
| 0.498 | 0.006 |  |  |  | + | 8 | -30.107 | 79.422 | 8.771 | 0.01 |
| 0.507 | 0.006 |  | -0.22 |  |  | 3 | -35.684 | 79.838 | 9.188 | 0.01 |
| 0.578 | 0.006 | -0.034 |  |  | + | 9 | -29.768 | 81.256 | 10.605 | 0.004 |
| 0.428 | 0.005 | 0.032 | -0.233 |  |  | 5 | -35.494 | 81.701 | 11.051 | 0.003 |
| 0.517 | 0.005 |  |  |  |  | 3 | -39.643 | 85.564 | 14.914 | 0 |
| 0.541 | 0.005 | -0.01 |  |  |  | 4 | -39.626 | 87.722 | 17.071 | 0 |
| 0.929 |  |  | -0.198 |  | + | 7 | -38.597 | 94.041 | 23.39 | 0 |
| 0.714 |  | 0.075 | -0.237 |  | + | 8 | -37.562 | 94.675 | 24.025 | 0 |
| 0.929 |  |  |  |  | + | 6 | -41.381 | 97.02 | 26.369 | 0 |
| 0.929 |  |  | -0.206 |  |  | 3 | -45.415 | 97.109 | 26.458 | 0 |
| 0.696 |  | 0.082 | -0.241 |  |  | 3 | -44.356 | 97.183 | 26.533 | 0 |
| 0.854 |  | 0.027 |  |  | + | 7 | -41.299 | 99.125 | 28.474 | 0 |
| 0.929 |  |  |  |  |  | 2 | -48.25 | 100.639 | 29.988 | 0 |
| 0.817 |  | 0.039 |  |  |  | 2 | -48.003 | 102.284 | 31.633 | 0 |

Table S7. Group levels (i.e., random effects) from a Bayesian hierarchical model predicting the daily survival rate of Hudsonian godwit chicks from 2009 ̶ 2019 near Beluga River, Alaska. Ȓ is the Gelman–Rubin statistic where Ȓ < 1.1 is evidence of convergence (Gelman & Rubin, 1992). Individual histories were grouped by study year, brood, and plot.

| Group level | Posterior Mean (SD) | 95% Credible  Interval | Effective  Sample Size | Ȓ |
| --- | --- | --- | --- | --- |
| 2009 | 0.05 (13.64) | -28.12, 30.87 | 14279 | 1.0 |
| 2010 | 0.01 (13.67) | -28.99, 29.99 | 15274 | 1.0 |
| 2011 | 0.1 (13.49) | -30.53, 27.12 | 15000 | 1.0 |
| 2014 | -0.15 (13.37) | -28.87, 27.64 | 15000 | 1.0 |
| 2015 | 0.05 (13.6) | -28.91, 28.86 | 15167 | 1.0 |
| 2016 | 0.08 (7.13) | -23.6, 2.56 | 14041 | 1.1 |
| 2019 | 0.11 (6.1) | -12.08, 11.35 | 14901 | 1.2 |
| 2009GN001 | 0.14 (8.86) | -17.3, 18.24 | 15000 | 1.0 |
| 2009GN002 | 0.06 (8.76) | -17.72, 17.84 | 16088 | 1.0 |
| 2009GN003 | -0.08 (8.67) | -17.92, 16.96 | 14798 | 1.0 |
| 2009GN007 | 0.11 (8.69) | -17.57, 17.56 | 16964 | 1.0 |
| 2009GN010 | -0.02 (8.73) | -17.27, 17.91 | 15639 | 1.0 |
| 2009GN012 | 0.11 (8.56) | -16.36, 18.09 | 14182 | 1.0 |
| 2009GN014 | 0.02 (8.76) | -18.02, 17.25 | 14961 | 1.0 |
| 2009GN018.2 | 0 (8.73) | -17.93, 17 | 14634 | 1.0 |
| 2009GN022 | -0.01 (8.65) | -18.94, 16.39 | 15000 | 1.0 |
| 2009GN027 | -0.01 (8.67) | -17.87, 16.76 | 14758 | 1.0 |
| 2009GN0282 | 0.08 (8.63) | -17.23, 17.5 | 15000 | 1.0 |
| 2009GN044 | -0.1 (8.71) | -17.64, 17.27 | 14861 | 1.0 |
| 2009GN045 | -0.1 (8.78) | -18.39, 16.86 | 16001 | 1.0 |
| 2009GN046 | -0.2 (8.7) | -18.62, 16.41 | 14546 | 1.0 |
| 2009GN047 | 0 (8.59) | -16.63, 17.95 | 15000 | 1.0 |
| 2009GN049 | 0.03 (8.62) | -17.17, 17.34 | 15000 | 1.0 |
| 2010GN11 | -0.02 (8.7) | -17.03, 17.74 | 14803 | 1.0 |
| 2010GN47 | 0.02 (8.81) | -18.13, 17.77 | 14507 | 1.0 |
| 2010GN58 | -0.06 (8.77) | -18.62, 17.25 | 14934 | 1.0 |
| 2010GN61 | -0.04 (8.75) | -18.04, 17.51 | 14517 | 1.0 |
| 2010GN62 | 0.02 (8.7) | -17.51, 17.35 | 14532 | 1.0 |
| 2010GN63 | -0.06 (8.7) | -17.69, 17.42 | 15466 | 1.0 |
| 2010GNGPM | -0.1 (8.59) | -18.01, 16.85 | 15080 | 1.0 |
| 2010GNHUYU | -0.01 (8.67) | -18.03, 17.33 | 15000 | 1.0 |
| 2010GNPE | -0.01 (8.74) | -17.29, 18.19 | 14097 | 1.0 |
| 2010GNUL | 0.01 (8.78) | -18.5, 16.97 | 15142 | 1.0 |
| 2010GNXEXY | 0.1 (8.63) | -16.61, 17.69 | 15418 | 1.0 |
| 2010GNYN2 | 0.07 (8.64) | -17.32, 17.49 | 16213 | 1.0 |
| 2010GNYTXL | -0.01 (8.51) | -18.03, 16.35 | 14345 | 1.0 |
| 2011GN13 | -0.01 (8.67) | -17.26, 17.53 | 15000 | 1.0 |
| 2011GNAPAU | -0.07 (8.74) | -18.48, 16.33 | 15215 | 1.0 |
| 2011GNC4T6 | -0.09 (8.7) | -17.62, 17.25 | 15352 | 1.0 |
| 2011GNC8J2 | 0 (8.71) | -17.68, 17.39 | 15627 | 1.0 |
| 2011GNCT | -0.03 (8.65) | -17.23, 17.63 | 14886 | 1.0 |
| 2011GNE5E9 | -0.01 (8.66) | -18.19, 16.62 | 16547 | 1.0 |
| 2011GNEAE7 | 0.02 (8.71) | -17.49, 17.98 | 15237 | 1.0 |
| 2011GNEE | -0.1 (8.68) | -17.61, 17.29 | 15149 | 1.0 |
| 2011GNH7T2 | -0.02 (8.79) | -18.41, 17.55 | 15870 | 1.0 |
| 2011GNH8LO | -0.02 (8.69) | -17.94, 17.17 | 15000 | 1.0 |
| 2011GNJ5K7 | -0.06 (8.63) | -17.96, 16.89 | 15812 | 1.0 |
| 2011GNJ6J0 | -0.01 (8.76) | -17.6, 17.41 | 15054 | 1.0 |
| 2011GNJTMU | -0.15 (8.72) | -17.57, 17.3 | 14835 | 1.0 |
| 2011GNK0PM | -0.02 (8.69) | -18.08, 16.74 | 15140 | 1.0 |
| 2011GNK4T7 | 0.04 (8.68) | -17.54, 17.8 | 14034 | 1.0 |
| 2011GNM2P2 | 0.06 (8.59) | -17.17, 17.36 | 15133 | 1.0 |
| 2011GNM3U0 | -0.05 (8.56) | -17.72, 16.9 | 14778 | 1.0 |
| 2011GNN8X3 | 0.05 (8.85) | -17.29, 18.38 | 14942 | 1.0 |
| 2011GNTANA | -0.1 (8.67) | -17.42, 17.53 | 15048 | 1.0 |
| 2011GNV9C0 | 0.1 (8.66) | -17.55, 17.17 | 15378 | 1.0 |
| 2011GNX50H | -0.04 (8.69) | -17.13, 17.56 | 17680 | 1.0 |
| 2011GNX6H3 | 0 (8.75) | -17.21, 18.3 | 15552 | 1.0 |
| 2011GNY9L6 | 0.05 (8.69) | -17.11, 17.55 | 14683 | 1.0 |
| 2014BHD17 | -0.09 (8.65) | -16.81, 18.31 | 15443 | 1.0 |
| 2014BHD19 | 0.06 (8.56) | -17.82, 16.69 | 15000 | 1.0 |
| 2014BJL17 | 0.03 (8.74) | -17.33, 17.81 | 14793 | 1.0 |
| 2014BJL18 | 0.08 (8.62) | -16.91, 17.5 | 15561 | 1.0 |
| 2014BJL19 | 0.01 (8.74) | -16.72, 18.86 | 14404 | 1.0 |
| 2014BJL25 | 0.12 (8.63) | -18.14, 16.59 | 15273 | 1.0 |
| 2014GJM06 | -0.11 (8.65) | -16.67, 18.44 | 15000 | 1.0 |
| 2015GJM05 | 0.08 (8.6) | -16.51, 18.2 | 15402 | 1.0 |
| 2015GJM18 | -0.03 (8.65) | -17.03, 18.23 | 15705 | 1.0 |
| 2015GJM35 | 0.09 (8.66) | -17.71, 17.12 | 15000 | 1.0 |
| 2015GJM36 | -0.02 (8.59) | -17.06, 17.39 | 14401 | 1.011 |
| 2015JAK05 | 0.04 (8.66) | -17.56, 17.46 | 14887 | 1.0 |
| 2015JAK21 | 0.01 (8.71) | -16.73, 18.05 | 16195 | 1.0 |
| 2015JMH10 | 0.01 (8.76) | -18.48, 16.96 | 15801 | 1.0 |
| 2015JMH20 | 0.1 (8.82) | -17.38, 18.13 | 15056 | 1.0 |
| 2015JMH28 | 0.01 (8.81) | -18.11, 17.84 | 16209 | 1.0 |
| 2015KJP18 | -0.01 (8.68) | -18.69, 16.75 | 14590 | 1.0 |
| 2015KJP44 | 0.08 (8.63) | -16.9, 17.92 | 15242 | 1.0 |
| 2015RJS05 | -0.03 (8.57) | -18.11, 16.63 | 14870 | 1.0 |
| 2015RJS05 | 0.01 (8.62) | -16.81, 17.7 | 15000 | 1.0 |
| 2015U1MUV | -0.08 (8.74) | -18.08, 17.02 | 15000 | 1.0 |
| 2016KRS48 | -9.52 (7.63) | -23.78, 7.31 | 12247 | 1.02 |
| 2016LKF04 | 0.56 (8.09) | -15.47, 16.61 | 13272 | 1.0 |
| 2016LKF22 | 0.01 (7.83) | -14.38, 15.86 | 15014 | 1.01 |
| 2016MLS14 | 0.93 (7.97) | -15.23, 16.6 | 13951 | 1.0 |
| 2016MLS37 | 1.58 (6.71) | -11.8, 15.43 | 12726 | 1.01 |
| 2016RIG15 | -0.33 (6.32) | -12.24, 13.79 | 12952 | 1.01 |
| 2016RJS04 | 0.4 (7.66) | -13.98, 16 | 12602 | 1.0 |
| 2016RJS07 | 0.16 (7.88) | -14.45, 16.73 | 12428 | 1.01 |
| 2016RJS10 | -3.46 (5.03) | -14.37, 6.13 | 12259 | 1.08 |
| 2016RJS16 | 3 (7.09) | -11.12, 17.81 | 12935 | 1.02 |
| 2019GB01 | -7.09 (3.08) | -13.5, -1.65 | 14744 | 1.01 |
| 2019GB02 | -2.31 (4.56) | -11.09, 7.26 | 14948 | 1.05 |
| 2019GB03 | -7.96 (4.41) | -15.49, 1.97 | 13044 | 1.07 |
| 2019GN01 | 1.6 (4.38) | -6.11, 11.44 | 15159 | 1.08 |
| 2019GN02 | 10.11 (5.97) | -0.66, 22.54 | 15057 | 1.02 |
| 2019GN03 | -8.88 (4.4) | -16.79, 0.63 | 13173 | 1.07 |
| 2019GN05 | 2.89 (6.6) | -7.96, 16.62 | 14273 | 1.04 |
| 2019GN06 | 8.15 (5.67) | -2.12, 19.74 | 14424 | 1.01 |
| 2019GN08 | 0.65 (2.99) | -5.79, 5.92 | 13055 | 1.01 |
| 2019GN09 | 6.69 (6.41) | -6.64, 19.79 | 12900 | 1.01 |
| 2019GN10 | -3.69 (6.79) | -15.05, 11.02 | 12128 | 1.01 |
| 2019GN11 | -5.76 (3.11) | -12.24, 0.12 | 13962 | 1.02 |
| 2019GN12 | 8.23 (5.62) | -1.68, 19.93 | 14157 | 1.01 |
| South Plot | 0.26 (4.43) | -8.25, 9.21 | 14071 | 1.16 |
| North Plot | 3.28 (4.78) | -3.97, 14.22 | 12493 | 1.02 |

1. Gelman, A., & Rubin, D. B. (1992). Inference from Iterative Simulation Using Multiple Sequences. Statistical Science, 7(4), 457–472. https://doi.org/10.1214/ss/1177011136

Table S8. Seasonal daily survival rates (DSR) of Hudsonian godwit chicks (*n* = 122) in Beluga River, Alaska among study years. DSR estimates from a Bayesian hierarchical model were extrapolated to 28 days as an estimate of percent fledged, and associated delta error is reported.

| Year | No.  chicks | DSR  (mean) | SD | 95% credible  interval | Estimated % fledged  (± delta error) |
| --- | --- | --- | --- | --- | --- |
| 2009 | 16 | 0.931 | 0.088 | 0.871, 0.964 | 22.11 (±19.13) |
| 2010 | 16 | 0.913 | 0.178 | 0.810, 0.963 | 14.75 (±36.69) |
| 2011 | 24 | 0.964 | 0.072 | 0.929, 0.982 | 45.84 (±21.22) |
| 2014 | 07 | 0.849 | 0.174 | 0.672, 0.939 | 3.18 (±1.88) |
| 2015 | 17 | 0.868 | 0.139 | 0.781, 0.924 | 5.16 (±3.00) |
| 2016 | 20 | 0.936 | 0.097 | 0.890, 0.963 | 24.71 (±28.95) |
| 2019 | 22 | 0.944 | 0.070 | 0.906, 0.967 | 29.79 (±21.54) |

Table S9. Model selection table comparing univariate linear models of population mismatch models predicting seasonal fledging rates in a population of Hudsonian godwits near Beluga River, Alaska from 2009-2019. AIC_c_ = Akaike’s Information Criterion corrected for small sample sizes.

| Model | logLikelihood | AIC_c_ | ΔAIC_c_ | Model  weight (*w_i_*) | R^2^ |
| --- | --- | --- | --- | --- | --- |
| Whole Demand | -25.14 | 64.27 | 0 | 0.43 | 0.55 |
| Difference in  peak dates | -25.31 | 64.63 | 0.35 | 0.36 | 0.48 |
| Peak Demand | -28.54 | 67.08 | 2.81 | 0.11 | 0.26 |
| Curve height | -26.62 | 67.23 | 2.96 | 0.10 | 0.25 |

Figure S1.

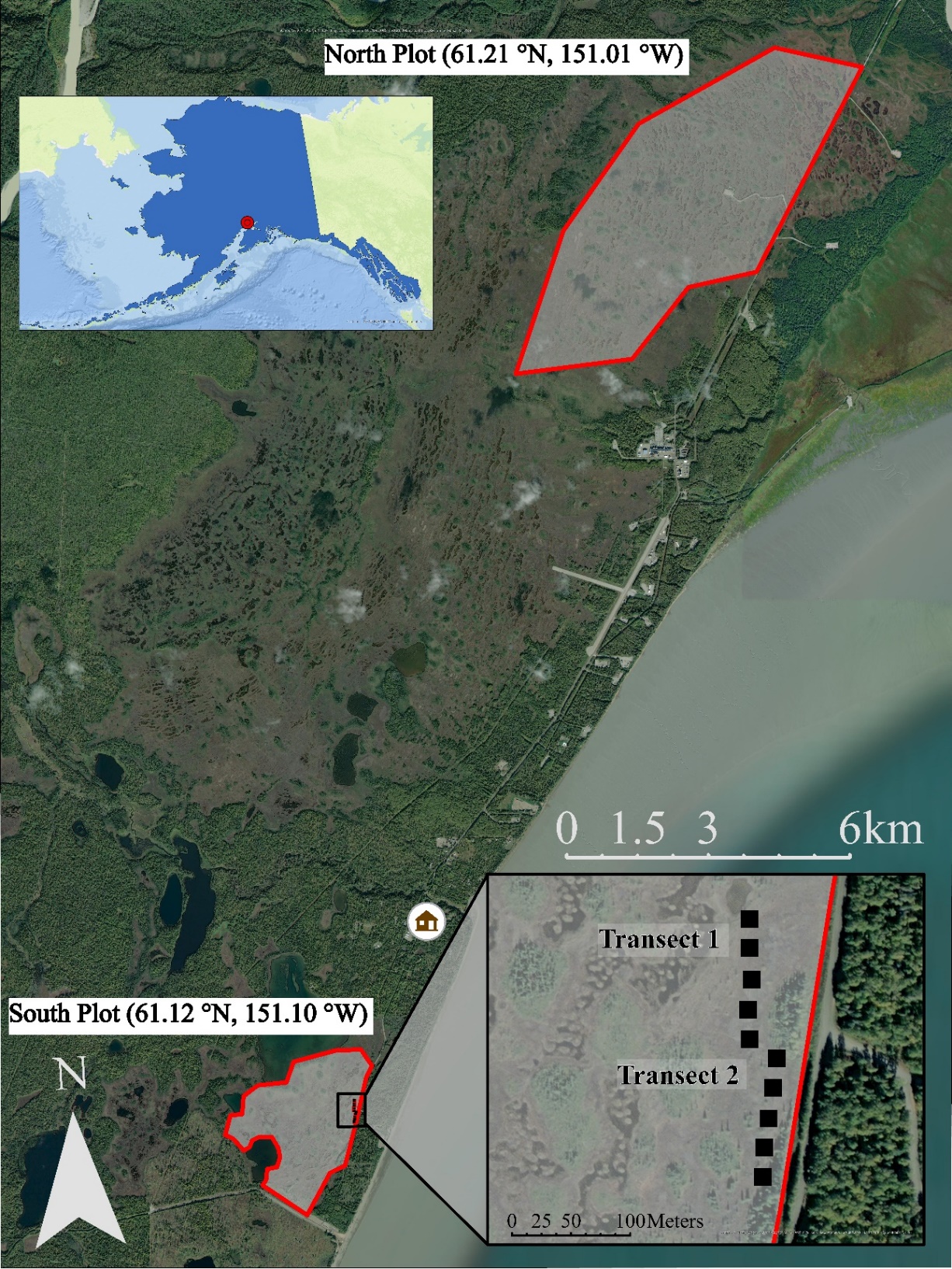

Figure S1. Map of North and South plots near the township of Beluga River, AK (house icon). (Inset top left) The study areas (red point) is located in southcentral Alaska on the west coast of the upper Cook Inlet. (Inset bottom right) The two, 100-m transects for daily invertebrate capture using pitfall traps (2009 ̶ 2011) or modified malaise traps (2014 ̶ 2016, 2019). Transects were placed according to Arctic Shorebird Demographic Network protocols (Brown et al. 2014).

Basemap images are the intellectual property of Esri and are used herein under license. Copyright © 2014 Esri and its Licensors. All rights reserved. Additional data sources: U.S. Census Bureau 2018.

Figure S2.

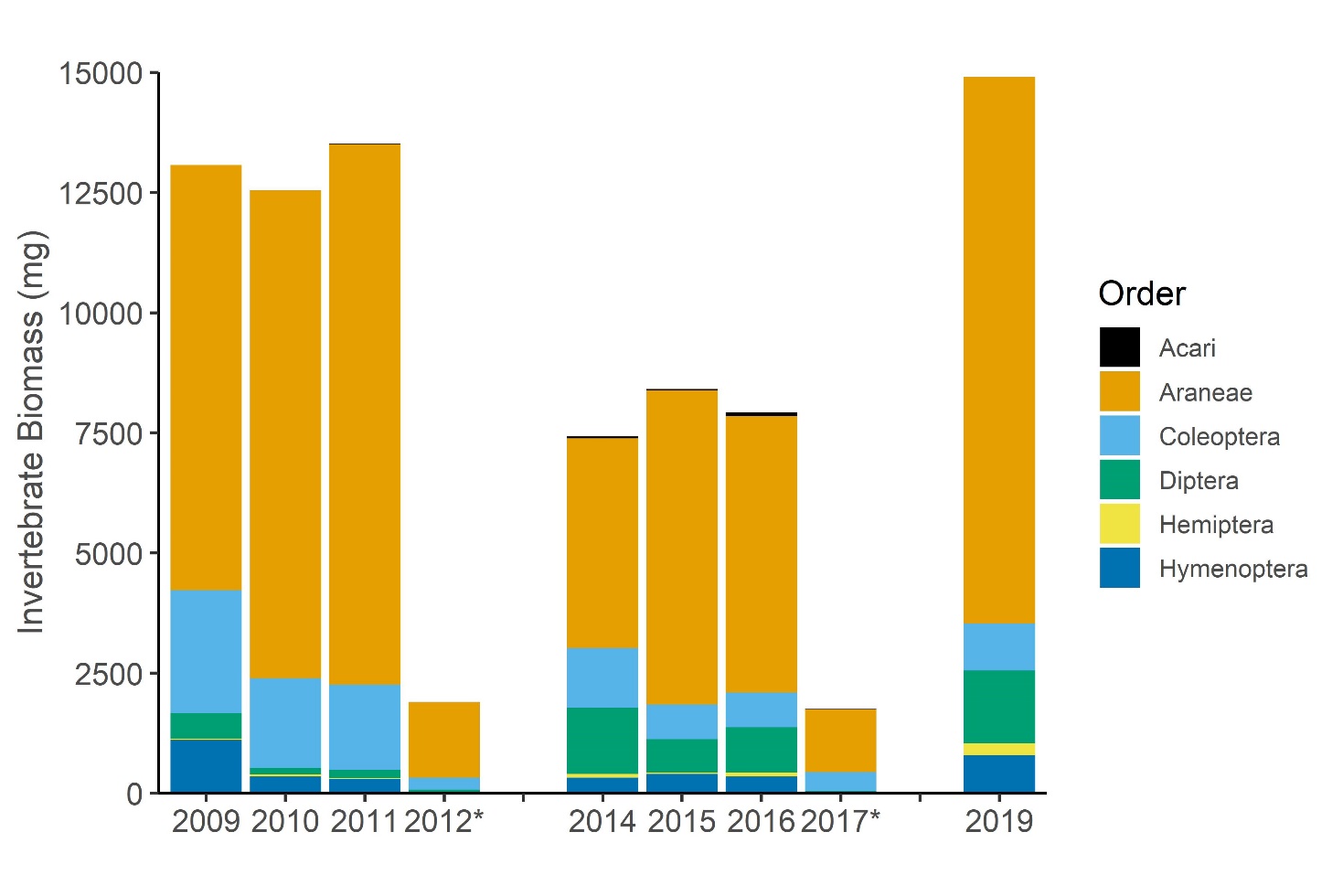

Figure S2. Interannual comparison of the available biomass and composition by each of the major Orders consumed by foraging godwit chicks. Invertebrates were monitored near Beluga River, AK from 2009 ̶ 2019. Biomass was determined using taxon-specific, length-weight relationships (Rogers et al., 1977; Ganihar, 1997; Robinson et al., 2018). The 2012 and 2017 seasons (*) were short seasons and do not represent the extent of the available energy. No monitoring occurred in 2013 and 2018.

1. Rogers, L. E., Buschbom, R. L., & Watson, C. R. (1977). Length-Weight Relationships of Shrub-Steppe Invertebrates. Annals of the Entomological Society of America, 70(1), 51-53. doi: 10.1093/aesa/70.1.51
2. Ganihar, S. R. (1997). Biomass estimates of terrestrial arthropods based on body length. Journal of Biosciences, 22(2), 219-224. doi: 10.1007/BF02704734
3. Robinson, S. I., McLaughlin, Ó. B., Marteinsdóttir, B., & O’Gorman, E. J. (2018). Soil temperature effects on the structure and diversity of plant and invertebrate communities in a natural warming experiment. Journal of Animal Ecology, 87(3), 634-646. doi: 10.1111/1365-2656.12798

Figure S3.

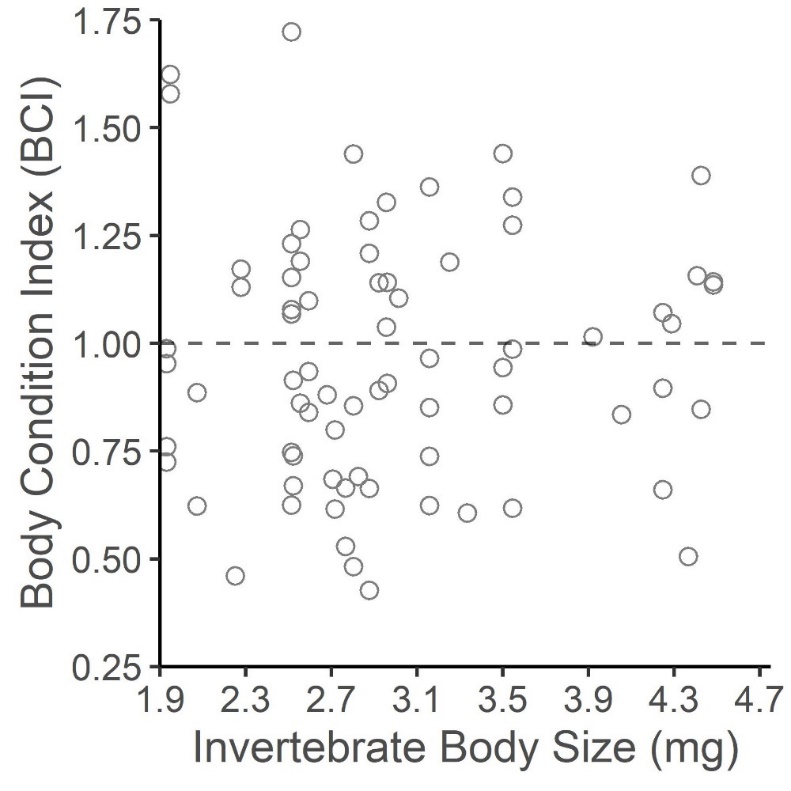

Figure S3. Effect of median invertebrate body size (mg) on the body condition index (BCI) of Hudsonian godwit chicks monitored near Beluga River, Alaska from 2009 - 2019. BCI is the ratio of observed to expected weight gain. BCI = 1 (dashed line) means individuals grew as expected, while BCI > 1 and BCI < 1 indicate better or worse than expected, respectively.

Figure S4.

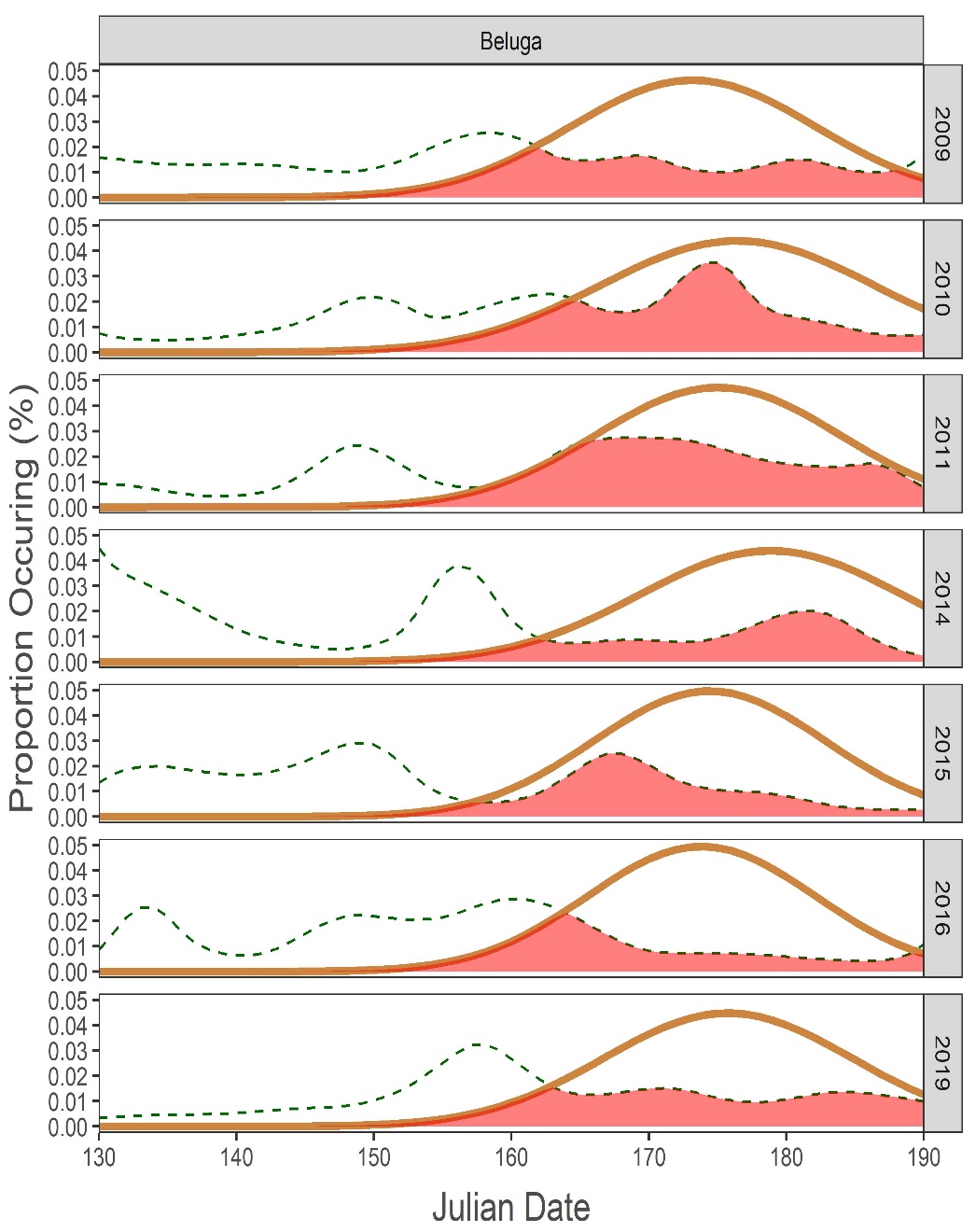

Figure S4. Overlap models of invertebrate biomass and Hudsonian godwit chick Whole Demand model (i.e., cumulative energetic requirements; Kilojoules d^-1^) for each year where both invertebrates and chicks were monitored between 2009 - 2019. Demand (red, solid line) and resource (black, dashed line) curves are represented as seasonal proportions, with the overlapping region (red, shaded) as a measure of ‘matching’. Each tile is a season.

Figure S5.

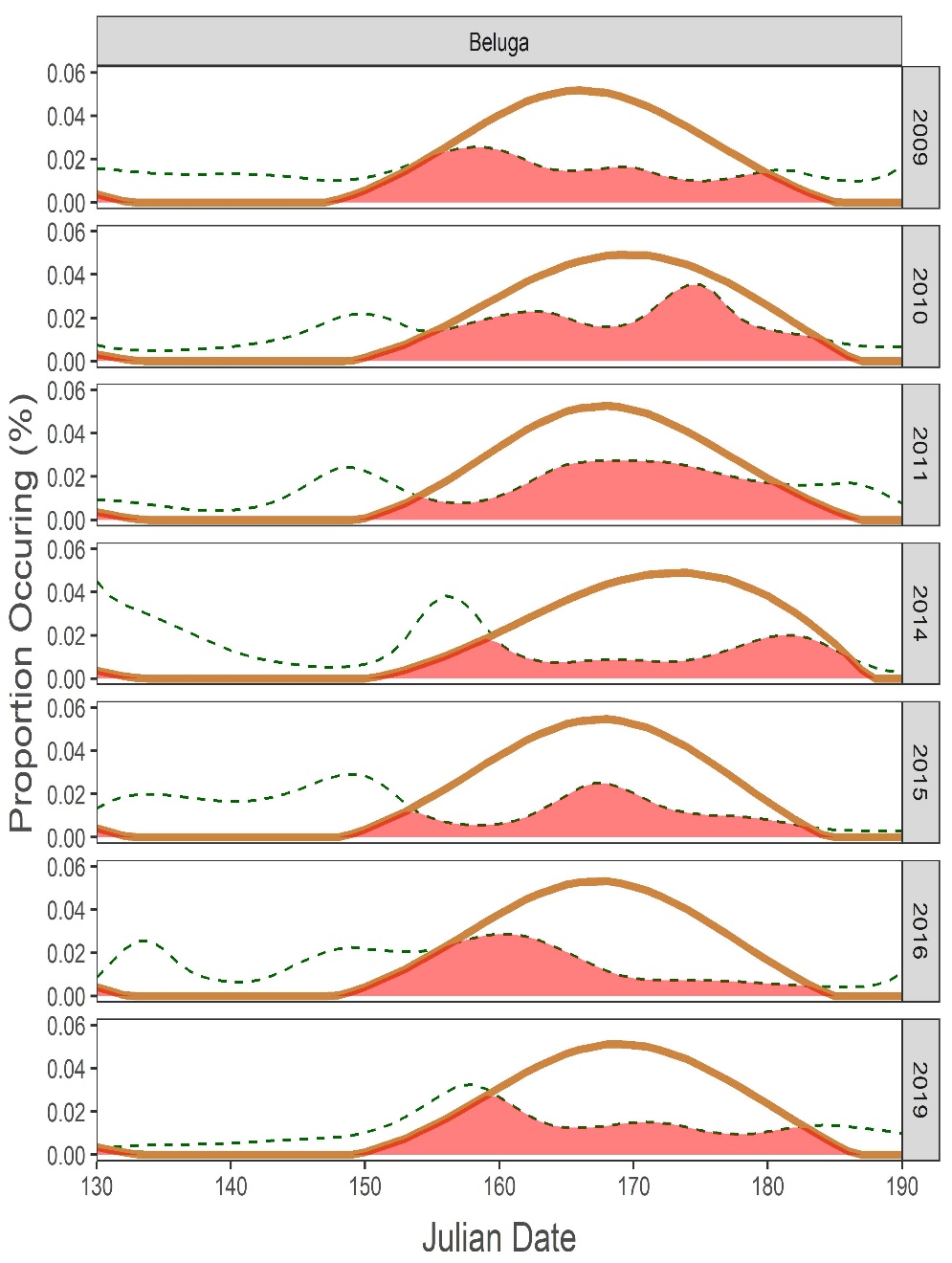

Figure S5. Overlap models of invertebrate biomass and Hudsonian godwit chick peak demand model (i.e., number of godwit chicks at age of peak growth rate per day) for each year where both invertebrates and chicks were monitored between 2009 - 2019. Demand (red, solid line) and resource (black, dashed line) curves are represented as seasonal proportions, with the overlapping region (red, shaded) as a measure of ‘matching’. Each tile is a season.
