## Appendix S2 for "The anatomy of a phenological mismatch: interacting consumer demand and resource characteristics determine the consequences of mismatching"

R Code S1. Computer code in JAGS (Just Another Gibbs Sampler v.4.1.0; Plummer, 2012) in the R programming environment (v4.0.3, R Core Team 2020) using using ‘runjags’ and ‘rjags’ packages (JAGS 4.1.0; Plummer, 2013; Denwood, 2016). This script is used to input and process data lists, run model, and generate outputs and convergence statistics.

```
#Define terms for run.jags model in R

#NH[i,j]: matrix of encounter history beginning and ending on #first and last day of the season

#insect[i,j]: matrix of daily invertebrate biomass estimates from #3-day average prior to each day

#size[i,j]: matrix of median invertebrate body size estimates from #3-day average prior to each day

#hatch[i]: vector of hatch date

#age[i,j]: matrix of chick age for each day of individual’s life

#plot[i]: vector of study plot (North = 2, South = 1)

#brood[i]: vector of brood ID

#year[i]: vector of study year (2009 = 1, 2010 = 2, ... 2019 = 7)


#
#

#script to impement model in R v3.6.0 or later


#load the packages
x <- c("runjags", "rjags", "arm"); lapply(x, require, character.only = T)

#
#

# ensures that all parameters would be output. Otherwise, runjags has a ceiling on the number of parameters  summarized
runjags.options(force.summary=T)

#
#

#define functions to calculate the last day alive, first day alive:
get.last <- function(m) max(which(m %in% c(0,1)))
get.first <- function(m) min(which(m %in% c(0,1)))

# build logit and inverse logit functions:
fn_logit <- function(X) log(X / (1 - X))
fn_invlogit <- function(X)	1/(1+exp(-(X)))

#
#


#
init <- Sys.time()	# record the start time for the example script


# identify first and last day each individual was alive
f <- as.numeric(apply(NH, 1, get.first))
l <- as.numeric(apply(NH, 1, get.last))	# last day that each nest was alive

#
#


#define MCMC simulation settings
adapt <- 600 #number of iterations to discard
bi <- 1000 #number of burn in, discarded before chains stabilize
ni <- 5000 #number of iterations to run
nt <- 3 #thinning factor
nc <- 3 #number of chains to run, my CORE i7 has 6 cores


#
#


# begin model in run.jags

cat("model {
 	for(i in 1:nnest){
 		for(j in f[i]:(l[i]-1)){
 			logit(phi[i,j]) <- intercept + w.a* eff_biomass*insect[i,j] + w.b*eff_size*size[i,j] + w.c*eff_size_age_interaction*age[i,j]*size[i,j] + w.d*eff_hatch*hatch[i,1] + w.e*eff_age*age[i,j] + alpha[plot[j]] + alpha1[brood[j]] + alpha2[year[j]]
```

```
 		}
 	}
# likelihood:
 	for(i in 1:nnest){
 		for(j in (f[i]+1):l[i]){
 			mu[i,j]<-phi[i,j-1]*NH[i,j-1]
 			NH[i,j]~dbern(mu[i,j])
 		}
 	}


#
#

#monitor loglik for WAIC calculation
	loglikcell[i,j] <- logdensity.cat(NH[i,j],mu[i,j])

#get row sum - for heirarchical model
	loglik[i] <- sum(loglikcell[i,])
#
#

# create model selection process using indicator-variable approach:

#hold variance constant
   tau.total ~ dgamma(3.29,7.8)
   K <- w.a + w.b + w.c + w.d + w.e
   tau.model <- K*tau.total

#weights priors
   w.a ~ dbern(0.5)
   w.b ~ dbern(0.5)
   w.c ~ dbern(0.5)
   w.d ~ dbern(0.5)
   w.e ~ dbern(0.5)

#priors are diffuse, intercept restricted close to zero

 	# parameter priors
 	  intercept~dnorm(0,0.01)
       eff_age~dnorm(0,25)
       eff_size~dnorm(0,25)
       eff_size_age_interaction~dnorm(0,25)
 	  eff_biomass~dnorm(0,25)
 	  eff_hatch~dnorm(0,25)

 	#random effect priors
   for(j in 1:n.plot){
     alpha[j]~dnorm(0,tau.plot) ###I(-2,2)
   		}
   #int.plot ~ dnorm(0,0.001) #initial intercept of plot effect
   tau.plot<-pow(sigma.plot, -2)
   sigma.plot~dunif(0,25)


   #
    for(j in 1:n.brood){
     alpha1[j]~dnorm(0,tau.brood) ###I(-98,98)
   		}
   #int.brood ~ dnorm(0,0.001)
   tau.brood<-pow(sigma.brood, -2)
   sigma.brood~dunif(0,25)

   #
    for(j in 1:n.year){
     alpha2[j]~dnorm(0,tau.year) ###I(-7,7)
   		}
   #int.year ~ dnorm(0,0.001)
   tau.year<-pow(sigma.year, -2)
 sigma.year~dunif(0,25)


 	# back-transform expected DSR and nest survival to fledging

 	dsr_mean <- 1/(1+exp(-(intercept)))
 	nsurv_mean <- dsr_mean^d


 }",fill=TRUE)
}
> sink()


#run the model with constant insect effect

#
#

global_BF <-run.jags( model="HUGO_match.model_global.txt", data=list(NH=NH,insect=zinsect,size=zsize,plot=plot,year=year,brood=brood,l=l,f=f,d=d,age=zage,hatch=hatch,nnest=nrow(NH),n.year=n.year,n.plot=n.plot,n.brood=n.brood),monitor=c("dsr_mean","nsurv_mean","eff_age","eff_size_age_interaction","eff_size"," eff_biomass",eff_hatch","intercept","w.a","w.b","w.c","w.d",”w.e","loglik"),inits=function(){list(intercept=rnorm(1,0,0.2),eff_bimoass=runif(1),eff_size_age_interaction=runif(1),eff_age=runif(1),eff_hatch=runif(1),alpha=runif(n.plot),eff_size=runif(1),alpha1=runif(n.brood),alpha2=runif(n.year))},thin=nt, n.chains=nc, burnin=bi,adapt=adapt,sample=ni,psrf.target=1.1, method='rjparallel')
```

1. Plummer, M. (2013). Package rjags: Bayesian graphical models using MCMC. Version 3.10.
2. R Core Team (2020) R: A Language and Environment for Statistical Computing. RFoundation for Statistical Computing, Vienna, Austria.
3. Denwood, M. J. 2016. runjags: An R Package Providing Interface Utilities, Model Templates, Parallel Computing Methods and Additional Distributions for MCMC Models in JAGS. Journal of Statistical Software, 71(9), 1-25. doi:10.18637/jss.v071.i09

R Code S2. Computer code written in R programming environment (v4.0.3, R Core Team 2020) to generate data frames from input sources and perform overlap analyses.

```
#### Set up ####
#create function to load and install (missing) packages
foo <- function(x){
  for( i in x ){
    #  require returns TRUE invisibly if it was able to load package
    if( ! require( i , character.only = TRUE ) ){
      #  If package was not able to be loaded then re-install
      install.packages( i , dependencies = TRUE )
      #  Load package after installing
      require( i , character.only = TRUE )
    }
  }
}

#load or install packages
foo( c("dplyr", "data.table", "mgcv", "sfsmisc", "ggplot2", "gamlss", "schoenberg", "lme4", "wesanderson", "MuMIn", "gridExtra", "grid", "visreg"))

#clean old objects from R
rm(list=ls(all=TRUE))


#### Develop dataset ####
# set working directory
setwd("C:/Users/14064/Dropbox/Chapter 2/Final Materials/Data/Overlap")

#read input data
# 1. resource biomass
biom <- read.csv("./population_model_biomass.csv") # single column vector of sum, daily biomass value (mg)
# 2. data vector of consumer hatch dates for each day of season
hatch_dates <- read.csv("./population_model_consumer.csv") # single column vector of sum, daily count of consumer
# 3. Daily RMR by age, in units of kJ/day
metabolism <- read.csv("./population_model_metab.csv") # single column vector of metabolic data


# season range
first <- 129 #first Julian day of season
last <- 190 #last Julian day of season
peak_demand <- 11 #in units of ontogeny (days)
Jdate <- c(first:last)
#define length of period in development
ontogeny <- c(1:(nrow(metabolism)-1))
Year <- c(2009,2010,2011,2014,2015,2016,2019) #study years
# study years

loop_object <- vector("list", length(Year))

for(i in 1:length(Year)){
  loop_object[[i]] <- rep(Year[i],length(Jdate))
}
Year <- as.data.frame(do.call(cbind, loop_object)); Year <- data.frame(stack(Year[1:ncol(Year)])); Year <- data.frame(Year[,1]); names(Year) = "Year"

# define peak demand record

temp <- as.data.frame(matrix(,nrow = peak_demand, ncol = 1)); names(temp) <- "cons_num";temp[is.na(temp)] = 0
peak_num <- rbind(temp,hatch_dates)
temp1 <- nrow(peak_num)-nrow(temp)
temp2 <- nrow(peak_num)
peak_num <- as.data.frame(peak_num[1:(nrow(peak_num)-peak_demand),]); names(peak_num) <- "peak_num"

#define list of sites
Site <- ("Beluga")

# compile into data frame
pop_data <- data.frame(Site, Year, Jdate, biom, hatch_dates, peak_num); head(pop_data)

#create function to calculate the whole demand per day
create_whole_mat <- function(x){
#generate multiple lags (ref at https://gist.github.com/drsimonj/2038ff9f9c67063f384f10fac95de566)
lags <- seq(length(ontogeny))
lag_names <- paste("lag", formatC(lags, width = nchar(max(lags)), flag = "0"),sep = "_")
lag_functions <- setNames(paste("dplyr::lag(., ", lags, ")"), lag_names)
df1 <- as.data.frame(x %>% mutate_at(vars(cons_num), funs_(lag_functions)))
df1

#rename columns to generic numbers
names(df1) = seq(ncol(df1))

#replace na's with 0
df1[is.na(df1)]=0

#preview
head(df1);tail(df1)

#create empty vector for storing loop outputs
loop_object <- vector("list", length(ontogeny))

#function to calculate daily kJ
for(i in 1:length(ontogeny)){
  x <- df1[,4+i]
  loop_object[[i]] <- x * metabolism[i,1]
}

#process vector into dataframe and rename
whole_demand <- as.data.frame(do.call(cbind, loop_object)); whole_num <- as.data.frame(rowSums(whole_demand[,c(1:ncol(whole_demand))])); names(whole_num) = "whole_num"


#bring into data frame
x$whole_num <- whole_num

}

#
whole_num <- as.data.frame(pop_data %>%
   group_by(Year) %>%
   do(data.frame(value=create_whole_mat(.))))

data <- cbind(pop_data,whole_num);data <- data[,c(1:6,8)];head(data)

#data[data == 0] <- NA; head(data)

biom_perc <- as.data.frame(data %>%
              group_by(Year) %>%
              mutate(freq = mean_biom / sum(mean_biom))); biom_perc <- biom_perc[8]; names(biom_perc) = "biom_perc"

whole_perc <- (as.data.frame(data %>%
              group_by(Year) %>%
              mutate(freq = whole_num / sum(whole_num)))); whole_perc <- whole_perc[8]; names(whole_perc) = "whole_perc"

peak_perc <- (as.data.frame(data %>%
              group_by(Year) %>%
              mutate(freq = peak_num / sum(peak_num)))); peak_perc <- peak_perc[8]; names(peak_perc) = "peak_perc"

raw <- cbind(pop_data, biom_perc, peak_perc, whole_perc); head(raw)

#### Begin Analysis ####

raw <- raw[,1:9]

min(raw$Jdate); max(raw$Jdate)
# First replace NA values for biomass with '0'
#raw[which(is.na(raw$mean_biomass)==TRUE),]$mean_biomass <- 0
#raw[which(is.na(raw$per_biomass)==TRUE),]$per_biomass <- 0


# Now, the dataset has field observed values for the biomass,
# which is the seasonal percentile of total biomass per sample divided by the number of days in the sampling interval
# (usually three days).
# We will fit a generalized additive model (GAM) (i.e., fit a smooth curve) to the data
# and extract daily biomass values from the fitted curve.

# We use a for-loop to repeat this process for invertebrate and each of 6 shorebird species
# for each site and year.

# First create the list of sites and shorebird species in the dataset
sitelist <- unique(as.character(raw$Site)) #only Beluga here
specieslist <- c('hugo') # only hugo here

# Two blank matrices to save outputs from the for-loops:
# 'output1' will have 10 columns to save the daily values from fitted curves
# 'output2' will have 4 columns to save the estimated overlap area size
output1 <- matrix(nr=0, ncol=5)
output2 <- matrix(nr=0, ncol=4)


# for-loop begins here
for(s in 1:length(sitelist)){
  raw_site <- subset(raw, raw$Site==sitelist[s]) #subset by site
  yearlist <- unique(as.numeric(as.character(raw_site$Year))) #for the given site, get the list of available years
  for(y in 1:length(yearlist)){ #loop through the years
    raw_sy <- subset(raw_site, raw_site$Year==yearlist[y]) #subset by year within site

    #
    #---------------------------------------->
    # ENTER VARIABLES HERE ------------------>
    #---------------------------------------->
    #
    # columns in raw_sy into separate objects for model fitting
    date <- raw_sy$Jdate
    biomass <- raw_sy$biom_perc
    hugo <- raw_sy$whole_perc

    #
    #Invert curve
    M3.gam <- gamlss(biomass ~ ps(date + I(date^2), df = 10, degree = 2), family = BEZI); summary(M3.gam) #fitting the GLM
    M3pred <- predict(M3.gam, se = TRUE, type = "response")      #predict daily values from the fitted model

    #you can check the GLM results with plots
    #p <- par(mfrow = c(2, 2), mar = c(5, 4, 1, 2))
    #plot(date, biomass, type = "p")
    #plot(M3, se = TRUE)
    #plot(date, biomass, type = "p")
    #I1 <- order(date)
    #lines(date[I1], M3pred$fit[I1], lty=1)
    #lines(date[I1], M3pred$fit[I1]+2*M3pred$se[I1],lty=2)
    #lines(date[I1], M3pred$fit[I1]-2*M3pred$se[I1],lty=2)


    #HUGO curve
    # Mhugo <- glm(hugo ~ date + I(date^2))
    Mhugo.gam <- gamlss(hugo ~ date + I(date^2), family =BEZI); summary(Mhugo.gam)
    Mhugopred <- predict(Mhugo.gam, se = TRUE, type = "response")


    # fitted results for the given site and year
    # saved into a data frame
    d <- data.frame(matrix(nr=length(date), ncol=0))
    d$site <- sitelist[s] #in the first column, repeat the site name
    d$year <- yearlist[y] #in the second column, repeat the year name
    d$x = date
    d$a <- M3pred$fit
    d$b <- Mhugopred$fit


    # replace negative values with '0', as negative percentage occurrence doesn't make sense
    d[,c(5:5)][d[,c(5:5)] < 0] <- 0

    # combine results from different sites and years together
    output1 <- rbind(output1, d) #rbind everything


    ### Now to get the overlap area coefficient!
    ### Area that spans from 0 to N on x-axis and from 0 to M on y-axis
    ### can be calculated as the sum of Y values over X.

    ### For two curves, then, the overlapped area is basically
    ### the sum of smaller Y values of the two curves across X.

    # So, get the smaller value of the two curves for each date/site/year
    if(sum(as.numeric(as.character(raw_sy$hugo_per)>0))) d$hugo <- pmin(d$a, d$b) else d$hugo <- 0

    # Again, replace any negative values with '0'
    d$hugo[d$hugo<0] <- 0

    # integrate areas under curves
    total.hugo <- integrate.xy(d$x, d$a) + integrate.xy(d$x, d$b) # this is the area under each of the two curves combined
    intersection.hugo <- integrate.xy(d$x, d$hugo)                # this is the area of overlap


    # calculate overlap coefficient
    # which is calculated as the overlapped area under two curves multiplied by 2 and divided by
    # the sum of areas under the two curves.
    overlap.hugo <- 2 * intersection.hugo / total.hugo

    # calculated values saved into a matrix
    output_sy2 <- matrix(NA, nr=1, ncol=4)
    output_sy2[,1] <- rep(sitelist[s],1) #in the first column, repeat the site name
    output_sy2[,2] <- rep(yearlist[y],1) #in the second column, repeat the year name
    output_sy2[,3] <- c('hugo')
    output_sy2[,4] <- c(overlap.hugo)

    output2 <- rbind(output2, output_sy2) #rbind everything

  }
}

output1 <- as.data.frame(output1)
output2 <- as.data.frame(output2)

colnames(output2) <- c("site","year","species","match")
colnames(output1)[3:5] <- c("date","invert","hugo")

#output2[which(output2$match %in% c("0","NaN")),]$match <- NA
output2$match <- round(as.numeric(as.character(as.factor(output2$match))),2)


output2[which(output2$species=="hugo"),]

# PLOT ############################################################

output1$site <- as.factor(output1$site)


ticks.warm <- data.frame (t = c(121,141,161,202), l = c(121,141,161,202)) ###change to match your julian dates


plot <- ggplot(output1, aes(x = date)) +   theme_bw() + xlab("Julian Date") +  ylab("Proportion Occuring (%)") +  geom_line(aes(y=invert), colour="darkgreen", linetype = "dashed") +   geom_line(aes(y=hugo), colour="tan3",lwd = 1.3, linetype = "solid") +   geom_area(aes(y = pmin(invert, hugo)), fill = 'red', alpha = 0.5) + scale_x_continuous(breaks=c(ticks.warm$t), labels=c(ticks.warm$l)) +  theme(panel.grid.minor = element_blank()) +   theme(panel.grid.major = element_blank()) +  theme(plot.title = element_text(size=14, hjust = 0)) +  facet_grid(year~site) +  geom_text(data=output2[which(output2$species=="hugo"),], aes(x=197, y=0.02, label=match), size=4, colour="black", inherit.aes=FALSE, parse=FALSE)+theme(legend.position = "left")+ylim(0,.08) + xlim(145,200) + theme(axis.text.x = element_text(colour = "grey30", size = 9),  axis.text.y = element_text(colour = "grey30", size = 9), axis.title.x = element_text(colour = "grey30", size = 14, vjust=-.5), axis.title.y = element_text(colour = "grey30", size = 14, vjust=1.5)) #+ geom_point(aes(x = date, y=per_biomass), data = raw)


plot
```
